# Mutant FUS perturbs m^6^A regulation by repressing ALKBH5, while restoring m^6^A levels alleviates ALS pathology *in vivo*

**DOI:** 10.1101/2025.11.26.690809

**Authors:** Gaia Di Timoteo, Adriano Setti, Francesca Margarita, Angelo D’Angelo, Andrea Giuliani, Martina C. Biagi, Michela Di Salvio, Tiziana Santini, Gianluca Cestra, Irene Bozzoni

## Abstract

Amyotrophic lateral sclerosis (ALS) is a neurodegenerative disorder marked by the progressive loss of motor neurons, where aberrant stress granule (SG) dynamics has emerged as a hallmark, especially in cases involving mutations in RNA-binding proteins such as FUS and TDP-43. We previously showed that m^6^A RNA modification is elevated in FUS-associated ALS and that restoring its physiological levels through METTL3 inhibition is sufficient to rescue the aberrant stress granule phenotypes in different human cellular models. Importantly, here we show that reducing m^6^A levels ameliorates locomotor defects and neuromuscular junction (NMJ) abnormalities in *Drosophila melanogaster* models of ALS expressing the human FUS^P525L^ mutant. Moreover, we demonstrate that the increase in m^6^A is attributable to reduced expression of the m^6^A eraser ALKBH5, resulting from impaired translation caused by direct FUS binding to its transcript. We further show that ALKBH5 downregulation contributes to RNA and protein alterations observed in FUS-associated ALS, driving the abnormal SG dynamics characteristic of the disease. This study supports the hypothesis that restoring correct m^6^A levels could serve as a potential therapeutic strategy for ALS.

## Introduction

Amyotrophic lateral sclerosis (ALS) is a neurodegenerative disease characterized by the progressive loss of motor neurons, which eventually leads to patient death ^1^. Aberrant stress granule (SG) dynamics have emerged as a hallmark of ALS, particularly in genetic contexts involving mutations in RNA-binding proteins such as FUS and TDP-43 ^2–7^. In particular, mutations in the nuclear-localization domain of FUS provoke its cytoplasmic delocalization and its recruitment into SG under stress conditions^8,9^. In previous studies, we demonstrated that ALS-associated mutations in FUS promote the formation of persistent SG with altered RNA and protein compositions and delayed dissolution; moreover, we showed that this phenomenon correlates with increased levels of N⁶-methyladenosine (m^6^A)^2,3^.

N6-methyladenosine (m^6^A) is an RNA modification known for its roles in RNA metabolism and numerous biological processes ^10^. We previously demonstrated that, although m^6^A plays only a limited part in SG physiology in normal conditions, its decrease can mitigate the aberrant features of FUS-associated ALS granules^2^. Interestingly, several studies have shown that in multiple ALS conditions, m^6^A levels are elevated compared with those observed under normal circumstances ^11,12–14^.

Notably, decreasing or increasing m^6^A levels was sufficient to rescue or exacerbate, respectively, the aberrant SG phenotypes observed in FUS-associated ALS cellular models^2^. Moreover, m^6^A up-regulation in wild-type cells mimicked the altered SG dynamics observed in ALS, underscoring m^6^A as a functional regulator of SG dynamics^2^.

However, although these findings establish a causal link between m^6^A abundance and SG dysfunction in cellular models of ALS, whether modulating m^6^A can slow disease progression *in vivo* remains unresolved. Also, the molecular mechanisms underlying m^6^A accumulation in FUS-ALS, and how this modification influences SG dynamics and composition, remained unclear.

*Drosophila melanogaster* has long served as a powerful model for investigating neurodegenerative diseases and conducting high-throughput screens for drugs and genetic modifiers. Notably, here we show that reducing m^6^A levels ameliorates locomotor defects and neuromuscular junction (NMJ) abnormalities in *Drosophila melanogaster* models of ALS carrying the human FUS^P525L^ mutation^15^, providing *in vivo* evidence of the beneficial effect of targeting m^6^A in ALS.

Furthermore, we show that elevated m^6^A levels in ALS cellular models arise from decreased expression of the m^6^A eraser ALKBH5. In particular, we demonstrate that FUS binds the ALKBH5 transcript, impairs its translation, and consequently lowers ALKBH5 protein abundance. Moreover, we found that ALKBH5 downregulation accounts for a subset of the RNA- and protein- alterations observed in FUS-associated ALS, and that some of these downstream changes drive the abnormal SG dynamics characteristic of the disease.

In summary, by demonstrating *in vivo* efficacy and revealing molecular insights upstream and downstream of m^6^A deregulation in ALS, our data not only further support m^6^A modulation as a therapeutic strategy for ALS but also identify additional factors that could be exploited therapeutically.

## Methods

### Fly strains, and treatments

Drosophila stocks and crosses were maintained on Drosophila standard medium (Nutri-fly, Genesee Scientific) at 25°C, unless otherwise indicated. RNAi fly lines targeting METTL3 expression and GAL4 drivers were obtained from Bloomington stock center (http://flybase.bio.indiana.edu/) and detailed as follows: 80431(y[1] sc[*] v[1] sev[21]; P{y[+t7.7]v[+t1.8]=TRiP.HMS06011}attP40); 80448(y[1] v[1]; P{y[+t7.7] v[+t1.8]=TRiP.HMS06028}attP2); 9146(w[1118]; (P{GMR-GAL4.w[-]}2/CyO); 1774(69B-GAL4 (w[*]; P{w[+mW.hs]=GawB}69B). Transgenic flies expressing hFUS^P525**L**^ were previously generated^15^.

### Climbing assay

Negative geotaxis assays were performed according to standard protocols. Adult male flies were separated and allowed to recover for 24 h to eliminate anesthesia effects. Flies were then transferred to empty vials and acclimated for 1 h before testing. Climbing ability was assessed by tapping flies to the bottom of the vial and recording the number of them reaching a 4 cm mark within 5 s. Flies were tested in groups of 10–15, with three trials conducted per assay.

### Immunofluorescence experiments in *D. Melanogaster*

Muscle filets from Drosophila third instar larvae were dissected according to standard protocols^13^. Briefly, larvae were dissected in PBS and fixed in 4% paraformaldehyde, 4% sucrose for 45’ at RT. Filets were permeabilized in PBST (0.25% Triton-X 100) 10’ at RT, and blocked 1 h at RT (PBS, 0.1 % Triton-X 100, 3% BSA), before to be incubated with primary Abs diluted in blocking buffer (PBS, 0.1 % Triton-X 100, 3% BSA) 1h at RT in a wet chamber. Samples were washed with PBS and incubated with secondary Abs or labeled dyes 1 h at RT. After 3 washes with PBS, preparations were mounted in antifade mounting medium with DAPI (Vectashield, Vector Laboratories, Newark, CA, USA). Confocal imaging was performed on LSM700 confocal laser scanning microscope (Zeiss, Oberkochen, Germany) equipped with four tunable light laser sources: 405, 488, 561, and 639 nm. Sequential confocal images from LSM700 were acquired using a 63x oil-immersion objective with a 1,024 × 1,024 format, z-step size of 0.5-1 μm, and with an electronic zoom magnification up to 1.0. Images were imported into ZEN 3.1 Blue Edition (Carl Zeiss Microscopy GmbH, Germany) software to obtain best fit maximum projections, which were analysed with NHI ImageJ FIJI freeware to evaluate differences of larval neuromuscular junction (NMJ) morphology in different genetic backgrounds. The maximum projections were then assembled on Adobe Photoshop CS4. Quantitative analysis of NMJ markers was performed on images with a manual method based on the delimitation of regions of interest by using NHI ImageJ FIJI freeware.

The following dyes and Abs were employed: anti horseradish peroxidase (HRP)-FITC #123-545-021 (Jackson ImmunoResearch, West Grove, PA, USA)(1:300); and fluorescently labeled phallodin Acti-stain™ 555 (Cytoskeleton, Denver, CO, USA) (1:300).

### Statistical analysis related to *D. Melanogaster* experiments

Data analyses were performed using the GraphPad Prism software 8.0.2 (GraphPad Software, San Diego, CA, USA). All experiments were independently repeated at least three times. Normality of sample distributions was assessed using the Shapiro–Wilk test prior to comparison. With normally distributed samples we used the unpaired t-test, otherwise Mann-Whitney test. For multiple comparisons, we used one way ANOVA and Tukey’s test. Each test was employed to evaluate the statistical significance that was set at P<0.05 and was reported by asterisks according to the following scheme: ***P<0.001, **P<0.01, and *P <0.05.

### Cell colture

SK-N-BE cells were cultured in RPMI (Sigma-Aldrich, Saint Louis, MO, USA) supplemented with 10% FBS (Sigma-Aldrich, #F2442), GlutaMAX supplement 1X (ThermoFisher Scientific, # 35050061), sodium-pyruvate 1mM (Thermo Fisher Scientific, #11360070) and Pen/Strep 1X (Sigma-Aldrich, #P4458). For the inducible expression of FUS-FLAG WT or P525L, SK-N-BE cells were exposed to 50 ng/mL Doxycycline (Sigma-Aldrich, #D9891) for 24h before the sodium arsenite treatment (Sigma-Aldrich, #106277). All cell lines were occasionally tested for mycoplasma contamination^46^.

For the CHX chase assay, 2.5x10^5^ cells were plated in 35mm plates of both FUS^WT^ and FUS^P525L^-espressing cell lines. After 24h each plate was exposed to 50 ng/mL Doxycycline (Sigma-Aldrich, #D9891) and after 24h treated with cycloheximide (100 µg/ml) or DMSO as control for 5 hours before harvesting^87^. For the experiments upon proteasome inhibition, 2.5x10^5^ cells were plated in 35mm plates of both FUS^WT^ and FUS^P525L^-espressing cell lines. After 24h each plate was exposed to 50 ng/mL Doxycycline (Sigma-Aldrich, #D9891) and after 24h treated with MG132 (50 µM) or DMSO as control for 5 hours before harvesting^87^.

### siRNA transfection

Reverse siRNA transfections were carried out according to manufactirer’s instruction by mixing 5 μl of Lipofectamine RNAiMAX Reagent (Thermo Fisher Scientific), siRNA (final concentration 30 nM) in 600 μl of Opti-MEM (Thermo Fisher Scientific), added with 800 × 10^3^ SK-N-BE cells diluted in 1ml of Opti-MEM (Thermo Fisher Scientific). 300 μl of the mixture was plated on collagen-coated glass coverslips in 24-well plates for stress granules counting experiments and the rest was plated in 35 mm plates for siRNA efficiency control experiments. The medium was replaced after 24 hours with fresh medium added with 50 ng/ml Doxycicline. After 24 hours the cells were treated with 0.5 mM of Sodium Arsenite for 1 hour. Control cells plates were harvested.

DS NC-1 (Integrated DNA Technologies, #51-01-14-04) was used as negative control. A complete list of the siRNA used for the RNA interference experiments are provided in table S1.

### Stress Granules Imaging and Quantification

SK-N-BE cells in the multiwell glass coverslips were fixed for 20 min at room temperature with cold 4% paraformaldehyde (Electron Microscopy Sciences, #15710) diluted in complete PBS (Sigma-Aldrich, #D1283), rinsed three times with complete PBS, and stored in PBS at 4 °C. Nuclei were stained with 1 μg/ml DAPI (#D9542, Sigma-Aldrich) in complete PBS for 5 min, and coverslips were mounted using ProLong™ Glass Antifade Mountant (Thermo Fischer Scientific, #P36980), leaving the slides on the bench overnight. Confocal images were acquired as Z-stacks (0.3-μm step size) using an inverted Olympus iX73 microscope equipped with an X-Light V3 spinning disc head (Crest Optics), a Prime BSI Scientific CMOS (sCMOS) camera (Photometrics), and MetaMorph software (Molecular Devices), with a 60× oil-immersion objective. Image analysis was performed using ImageJ software. GFP-G3BP1 served as a SG marker, and the ImageJ tool “3D Object Counter” was employed to quantify the number of SGs.

### RNA analyses

Total RNA was extracted with Direct-zol RNA Miniprep (Zymo Research) kit according to the manufacturer’s specifications. For immunoprecipitation experiments, the RNA was recovered through standard phenol-chloroform extraction and precipitation. When needed, DNAse I treatment was performed (Thermo Fisher Scientific). Reverse transcription reactions for routine experiments were performed using PrimeScript RT Master Mix (Takara Bio), while for RNA derived from IP experiments the SuperScript VILO cDNA Synthesis Kit (ThermoFisher Scientific) was used, according to manufacturer’s protocol. RT-qPCR analyses were performed using PowerUp SYBR Green Master Mix reagent (ThermoFisher Scientific), according to the manufacturer’s instructions. DNA amplification was monitored on an Applied BiosystemsTM 7500 Fast or StepOnePlus System qPCR instrument with the 7500 Software (Applied Biosystems) version 2.3 or with the StepOneTM Software (Applied Biosystems) version 2.3, respectively. Relative RNA quantity was calculated as the fold change (2−ΔΔCt) with respect to the experimental control sample set as 1 and normalized over GAPDH or ATP5O mRNA. A complete list of the oligonucleotides used for qRT-PCR experiments are provided in table S1.

### Protein analyses

Cells were harvested in a suitable volume of Protein Extraction Buffer (Tris pH 7.5 100 mM, EDTA 1 mM, SDS 2%), PIC1X (Complete, EDTA free, Roche) and incubated 10 min on ice, then incubated on a rotator for 20 min at 4 °C and centrifuged at 16000 × g for 10 min at 4 °C. The supernatant was transferred to a clean tube, used for subsequent analyses, or stored at −20 °C. Total protein concentration was measured through the Bradford reagent (Bio-Rad Protein Assay) following manufacturer’s instructions. 10-20 μg of proteins were loaded on 4–12% bis-tris polyacrylamide gel (Thermo Fisher Scientific) and transferred to a nitrocellulose membrane. The membrane was blocked in 5% milk and then hybridized with specific antibodies for 1 hr at room temperature or overnight at 4 °C. After three washes in TBST, the filter was hybridized with the corresponding secondary antibody, if required, for one hour at room temperature. All antibodies used in this study are reported below. Protein detection was carried out with WesternBright® ECL Chemiluminescent HRP Substrate (Advansta) or with Clarity Max Western ECL Substrate (Bio-Rad). Images were acquired using a ChemiDocTM MP Imager (Bio-Rad) and images were analyzed using Image LabTM 5.2.1 Software (Bio-Rad)^46^. The following antibodies were used in Western blots for protein analyses:

− Rabbit anti-METTL3 monoclonal antibody (Abcam, #ab195352, 1:1000)

− Rabbit anti-METTL14 polyclonal antibody (Atlas, #HPA038002, 1:1000)

− Rabbit anti-ALKBH5 polyclonal antibody ABclonal A11684 1:1000

− Rabbit anti-YTHDF1 polyclonal antibody ThermoFischer A305-850A-M Scientific 1:1000

− Rabbit anti-YTHDF2 polyclonal antibody Sigma-Aldrich ABE542 1:1000

− Rabbit anti-YTHDF3 polyclonal antibody Proteintech 25537-1-AP 1:1000

− Rabbit anti-YTHDC1 polyclonal antibody (1:500 Abcam ab122340)

− Rabbit anti-METTL16 polyclonal antibody (1:1000 Abcam ab185990)

− Rabbit anti-MAT2A polyclonal antibody (1:500 Abcam ab122340)

− Mouse anti-WTAP (D-7) monoclonal antibody Santa Cruz Biotechnology sc-374280 1:1000

− Rabbit anti-ACSL3 monoclonal antibody ABclonal A22085 1:10000

− Rabbit anti-HSPA1L polyclonal antibody ABclonal A1856 1:1000

− Rabbit anti-CAPN2 monoclonal antibody Santa Cruz Biotechnology sc-373966

− Rabbit anti-HSP70 monoclonal antibody Santa Cruz Biotechnology sc-32239 1:1000

− Rabbit anti-PPM1E monoclonal antibody ABclonal A9374 1:1000

− Rabbit anti-VGF polyclonal antibody Antibodies.com A96425 1:1000

− Rabbit anti-DYNLT3 polyclonal antibody ABclonal A16982 1:2000

− Rabbit anti-TPPP3 polyclonal antibody ABclonal A6775 1:1000

− Rabbit anti-KIF5C polyclonal antibody Proteintech 25897-1-AP 1:2000

− Rabbit anti-CRABP1 monoclonal antibody ABclonal A5434 1:1000

− Rabbit anti-DNM1 polyclonal antibody ABclonal A18563 1:2000

− Rabbit anti-CIRBP polyclonal antibody ABclonal A6080 1:1000

− Rabbit anti-MTMR9 polyclonal antibody ABclonal A13124 1:1000

− Rabbit anti-Ezrin monoclonal antibody ABclonal A19048 1:2000

− Rabbit anti-GIT1 polyclonal antibody ABclonal A15437 1:2000

− Anti-Flag M2-Peroxidase (HRP) Sigma-Aldrich A8592 1:2500

− Anti-ACTB-Peroxidase (AC-15) monoclonal antibody Sigma-Aldrich A3854 1:10000

− Rabbit anti-GAPDH (6C5) monoclonal antibody Santa Cruz Biotechnology sc-32233 1:10000

− Mouse anti-ACTININ (H-2) monoclonal antibody Santa Cruz Biotechnology sc-17829 1:10000

− Anti-GIOTTO was a gift from by Maria Grazia Giansanti (CNR Cat# Giotto_001, RRID:AB 2892585) 1:10000

− Anti-Rabbit IgG (H+L) Secondary Antibody, HRP Thermo Fisher Scientific 31460 1:10000

− Anti-Mouse IgG (H+L) Secondary Antibody, HRP Thermo Fisher Scientific 32430 1:10000

### CLIP assay

100 mm plates with SK-N-BE cells carrying FUS^P525L^ at maximum 80% confluency were washed twice with ice-cold PBS 1× (Sigma- Aldrich) and irradiated with 0.4J/ cm2 of 254 nm UV light. Cells were lysed in RIPA buffer (Tris-HCl pH 8 20mM, NaCl 100mM, EDTA 0.5mM, NP-40 0.5%, SDS 0,1%) supplemented with PIC 1× and RNase Inhibitor (Thermo Fisher Scientific). Lysates were incubated on ice for 15 min at 4°C, then passed through a 21 G needle. Lysates were spun down at 16000×g for 10min at 4°C and the supernatants were collected, then quantified with Bradford assay. 10% of the lysate was saved to be used as input, while for each immunoprecipitation 1 mg of extract was incubated with 30 μl of anti-FLAG beads (Sigma, # M8823) for 2 hours on a rotator at 4°C. Bead-bound antibody-RNA complexes were recovered on a magnetic rack, washed three times on a rotator for 2min at room temperature with 500μl Wash Buffer (Tris-HCl pH 7.4 50mM, NaCl 150mM, MgCl2 1mM, NP-40 0.05%) and three times with High-Salt Wash Buffer (Tris-HCl pH 7.4 50mM, NaCl 500mM, MgCl2 1mM, NP400.05%).One fifth was used for protein analysis, while 4/5 were used for RNA analysis after 1hr treatment with 10 μl Proteinase K (Roche) at 70°C in 90 μl PNK buffer (Tris-HClpH7.410mM, NaCl 100mM, EDTA 1mM, SDS 0.5%). After reverse-transcription of the extracted RNA with VILO cDNA Synthesis Kit (ThermoFisher Scientific), RT-qPCR was performed to evaluate targets enrichment^19^.

### RIP assay

10–15 Drosophila heads were homogenized in 400 µl RIPA (Tris-HCl pH 8 20mM, NaCl 100mM, EDTA 0.5mM, NP-40 0.5%, SDS 0,1%) supplemented with PIC 1× and RNase Inhibitor (Thermo Fisher Scientific), and left on ice for 15 min. Lysates were spun down for 15min at 4°C at 13000 rpm and the supernatants were collected. 200-500ug of extract quantified through a Bradford assay and diluted in up to 1 ml of IP buffer ((20mM TrisHCl pH7.5, 150mM NaCl, 15mM MgCl2, 0.5% NP40, 1mM EDTA, DTT 1 mM, PIC (Roche) 1x) and incubated with 10 µg of tRNA and 30µL Anti-FLAG Magnetic Beads (Sigma M8823) and incubated 2hrs at 4°C on a wheel. The 10% of the used extract was saved to be used as input. The beads were recovered through a magnetic rack and washed four times with 1ml wash buffer1 (20mM TrisHCl pH7.5, 200mM NaCl, 15mM, MgCl2, 0.5% NP40, 1mM EDTA). Samples collected after the last wash were divided for protein or RNA analysis. The proteins fraction was resuspended in LDS 1X buffer (Biorad) added with DTT 1µM, heated at 95°C for 10 min and loaded for western blot analyses. The RNA was extracted and reverse transcribed with VILO Superscript (Life Technologies) for qPCR analyses.

### FUS^P525L^ HITS-CLIP Analysis

RNA libraries for all samples were produced using Stranded Total RNA Prep with Ribo-Zero Plus (Illumina). All samples were sequenced on an Illumina Novaseq 6000 Sequencing system with an average of about 50 million 100 nt long paired-end read pairs.

The first Control replicate and both replicates of the mAID condition in this FUS^P525L^ HITS-CLIP study are included in this work, whereas the second biological replicate of the Control FUSP525L dataset was obtained from accession GSE242771. All samples: those presented here and the Control replicate from accession GSE242771 were produced within the same experimental batch.

For each condition (CTRL and mAID) peak calling and identification of reproducible peaks were performed as described in Mariani *et al.*, 2024 ^3^. The count matrix was generated by quantifying, for each BAM file from each sample, reads overlapping reproducible peaks using BEDtools intersect version 2.29.1^16^ considering only the start of read 2 as the reference for each fragment. To facilitate accurate data normalization and improve dispersion estimates, a background consisting of 200-nt genomic windows was also included. To define regions with differential FUS^P525L^ binding between two conditions, results from two complementary analyses were combined. The first analysis identified peaks enriched in the IP relative to the INPUT, consistently with the peak calling (see Materials and Methods), but using edgeR version 3.34.1^17^ with the following thresholds: log2FE(*IP/INPUT*) > 1, FDR < 0 and FPKMs > 5 in both control and METTL3 KD. The second analysis accounts for the identifications of regions displaying significant differences in enrichment between two tested conditions. This analysis makes use of TMM normalization and compares two conditions using the contrast formula:*(IP1−INPUT1)−(IP2−INPUT2)*, defining regions with specific enrichment for each condition (PValue < 0.05). A generalized linear model (GLM) approach was applied to both analyses, incorporating a blocked design to account for replicate variability, improve dispersion estimates, and minimize the influence of outliers. Peaks were classified as differentially enriched in one condition versus the other if they showed significant differential enrichment in the contrast while also maintaining consistent enrichment in the IP versus INPUT comparison; regions not meeting these criteria were considered invariant. Downstream visualization and identifications of differentially enriched peaks were performed using Python. Scatter density plots of log fold-changes were generated using JointGrid with KDE-smoothed marginal distributions

### RBP binding propensity analysis

To investigate whether additional RBPs could bind to the same region of ALKBH5 mRNA targeted by FUS^P525L^, we performed an RBP binding propensity analysis using in-vitro and eCLIP-based protein affinity metrics. In-vitro protein affinity was derived from RNAcompete experiments (Ray *et al.*, 2022) and retrieved from CISBP-RNA database^18^ while the eCLIP-based approach took advantage of PEKA scores^19^. For the analysis we used as reference the longest functional isoform of ALKBH5 gene (ENST00000399138) according to the Ensembl database (release 99). Transcript sequence scanned with sliding windows of 5 nt for PEKA scores and 7 nt for RNAcompete, depending on how the protein affinity scores were provided (5-mers and 7-mers, respectively). For each protein the affinity scores were mapped along the ALKBH5 transcript sequence based on k-mer windows and then normalized using Z-score. Furthermore, we employed BEDtools intersect version 2.29.1^16^ to screen POSTAR3 CLIPdb^20^ in order to identify other RBPs candidates having a coverage higher than 50 targeting the FUS binding region. Candidate RBPs were defined as those with strong affinity (Z-score > 3) for the FUSP525L binding region in the 3′UTR (nucleotides 2005–2205, as identified by FUSP525L HITS-CLIP).

### M^6^A CLIP

m^6^A CLIP was performed as previously described ^21,22^. Briefly, total DNase I-treated RNA was purified from SK-N-BE cells, 20 μg of RNA were diluted in IP buffer supplemented with RNase Inhibitor (Thermo Fisher Scientific) and incubated with 3 μg of anti-m^6^A antibody or IgG for 2 hrs at 4 °C rotating head over tail and crosslinked, 10% of the solution was saved to be used as input, the leftover incubated with protein A/protein G Dynabeads (Thermo Fisher Scientific) for 2 hrs at 4 °C. Bead-bound antibody-RNA complexes were washed and recovered. After phenol-chloroform extraction and precipitation, RNA was resuspended in 30 μl, and 7 μl were reverse-transcribed with VILO cDNA Synthesis Kit (ThermoFisher Scientific) in a 10 μl reaction. RT-qPCR was performed to evaluate targets enrichment.

## Results

### METTL3 KD relieves FUS-associated locomotion defects in flies

Given the promising results of our previous studies, particularly the beneficial effects of restoring physiological levels of m^6^A methylation in cellular models of FUS-associated ALS^2^, we aimed to validate these findings in an ALS *in vivo* animal model.

To assess the *in vivo* role of METTL3 on FUS-mediated toxicity, we exploited the UAS-GAL4 system^23^ to achieve a RNAi-mediated downregulation of the *Drosophila* ortholog of human METTL3 (IME4), in flies expressing the pathological human FUS^P525L^. Firstly, we assessed the efficiency of silencing *Drosophila* METTL3, which shares 45% identity and 58% of similarity with the human protein (DIOPT Ortholog Finder^24^), by expressing different RNAi constructs (#80450, #80431 and #80448, BDSC) (Fig. S1A) under the control of the eye specific GAL4 driver GMR. Downregulation of METTL3 was similar in all the strains tested (30-40%, Fig. S1A). We selected one of such strain showing consistent reduction of METTL3 mRNA (#80448, BDSC; hereafter named UAS-METTL3^RNAi^) for subsequent experiments. Thus, we used the 69B-GAL4 driver, which expresses

GAL4 panneuronally, in the embryonic epidermis and imaginal discs to guide the expression of UAS-METTL3-RNAi and UAS-hFUS^P525L^. We then tested whether the expression of hFUS^P525L^ increased the overall levels of m^6^A, as previously shown in human ALS cellular models^2^, and whether upon METTL3 knock down there was a concomitant decrease of m^6^A mRNAs. Through the colorimetric EpiQuik assay we confirmed both features in transgenic Drosophila extracts (Fig. S1B).

Although the increase in m^6^A levels was modest and slightly lower than in mammalian cell systems, it was consistent and reproducible; furthermore, because we used panneuronal expression of hFUS^P525L^, it is possible that our analysis, where we used whole heads, may have some non-neuronal tissue contamination.

We used these strains to assess the locomotor behaviour of these flies. To this aim, we performed a negative geotaxis (climbing) assay, which quantitatively measures the ability of flies to climb against gravity^25^. Flies expressing UAS-hFUS^P525L^ displayed a strong locomotor defect compared to controls, with a significantly lower percentage of animals reaching the target height (Fig. 1A, Fig. S1C). Notably, this strong impairment of locomotor ability observed in hFUS^P525L^ flies was significantly ameliorated by the downregulation of METTL3 (Fig. 1A).

**Figure 1.**
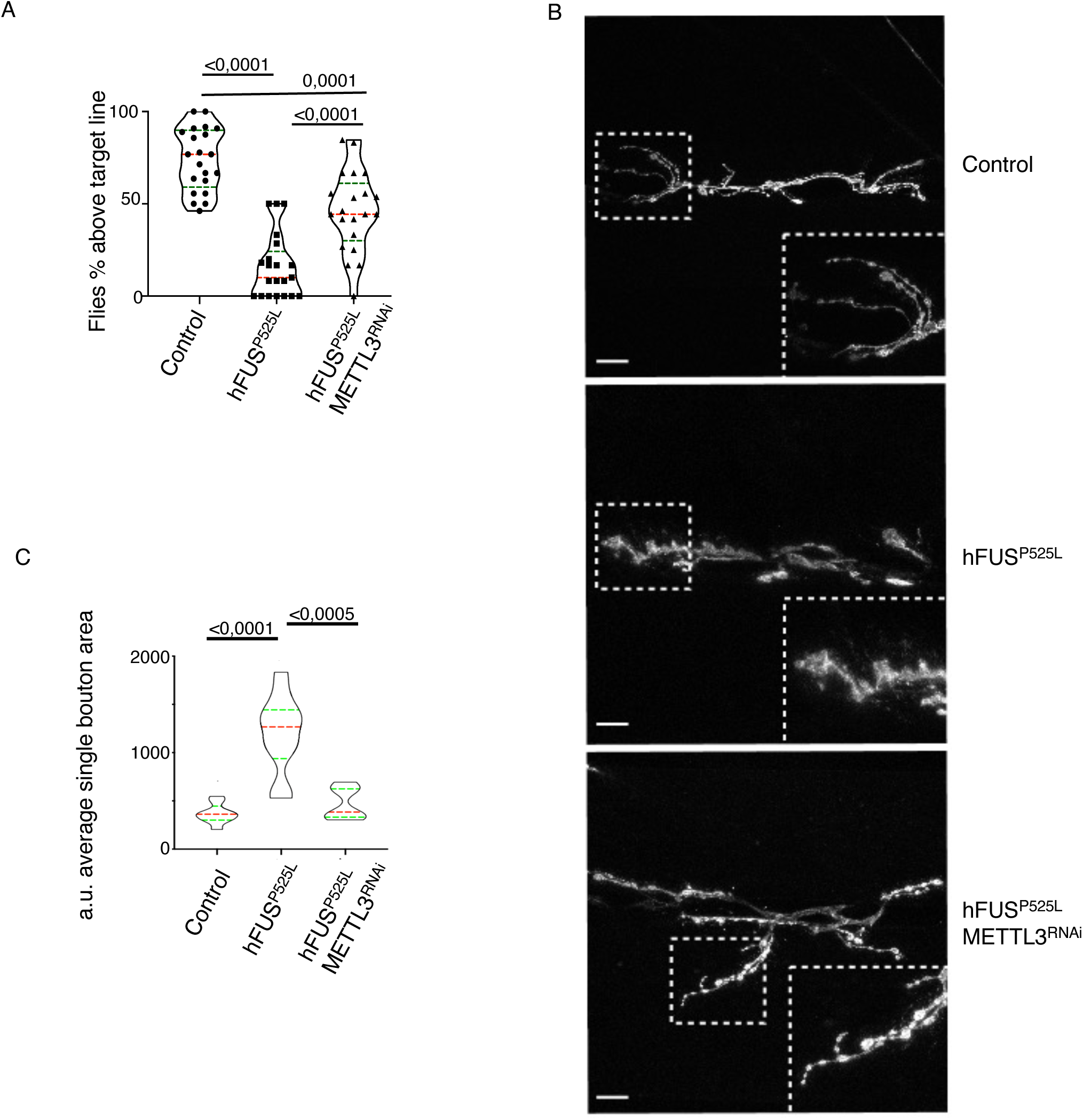
A) Quantification of fly locomotor activity assessing the percentage of adult male flies that express the indicated RNAi constructs or transgenes, under control of 69B-GAL4, and reach 4 cm in 5 seconds (control n=76 flies, hFUS^P525L^ n=77 flies, hFUS^P525L^/ METTL3^RNAi^ n=79 flies). The truncated violin plot reports median (dashed lines), first and third quartile (dotted lines) and density plot (outside lines). One way ANOVA and Tukey’s multiple comparison tests were used to compare samples, p-value are indicated. B) NMJs on muscle 6 and 7 of third instar larva expressing the indicated constructs under control of 69B-GAL4 driver were analyzed by IF. Representative images of NMJs labeled with anti-HRP to highlight presynaptic membranes. C) Bouton area quantified for each NMJ and averaged (Control n=210 boutons, hFUS^P525L^ n=153 boutons, hFUS^P525L^ METTL3^RNAi^ n=211 boutons) (control n=11 NMJs, hFUS^P525L^ n=11 NMJs, hFUS^P525L^METTL3RNAi n=7 NMJs). All crosses for the NMJ analysis were held at 29°C. Scale bars, 20mm. The truncated violin plots report median (dashed lines), first and third quartile (dotted lines) and density plot (outside lines). One way ANOVA and Tukey’s multiple comparison tests were used to compare samples, p-value are indicated.

Following on the impact of METTL3 downregulation on locomotor defects due to hFUS^P525L^, we assessed its effect on the morphology of larval neuromuscular junctions (NMJ) of muscles 6 and 7^26^. NMJ from third instar larvae expressing either UAS-hFUS^P525L^ alone or in combination with UAS-METTL3^RNAi^, were labelled with anti HRP, which stains the presynaptic neuronal membrane, to study the synaptic bouton morphology. In flies expressing UAS-hFUS^P525L^, we observed prominent alterations of bouton morphology (Fig. 1B) with a significant increase in the bouton average area compared to controls (Fig. 1C). Interestingly, this phenotype was significantly recovered upon METTL3 downregulation (Fig. 1C).

In conclusion, these data indicate that METTL3 downregulation is beneficial in ALS model flies both in terms of locomotion activity as well as for NMJ morphology.

### m^6^A alterations does not affect FUS binding

To investigate whether METTL3 knock-down affects the repertoire of FUS^P525L^ RNA interactors, we performed HITS-CLIP (as described in Di Timoteo et al., 2024^2^) in FUS^P525L^ SK-N-BE cells under control and METTL3 knock-down conditions (Table S2). To assess whether m^6^A depletion affects FUS^P525L^ binding, we performed differential binding analysis comparing control *versus* METTL3-KD samples. The analysis revealed only a very limited number of differential peaks (5 specific to the Control and 6 specific to the METTL3 knock-down condition; Fig. 2A, Fig. S2A, S2B and Table S2) corresponding to 5 interactors that lose and 6 that gain FUS ^P525L^ binding upon METTL3 depletion (0.48% and 0.62% of the 1027 total interactors detected, respectively; Fig. 2B). Altogether, these results indicate that the reduction of m^6^A levels following METTL3 knock-down does not substantially alter the repertoire of RNA targets interacting with FUS ^P525L^.

**Figure 2.**
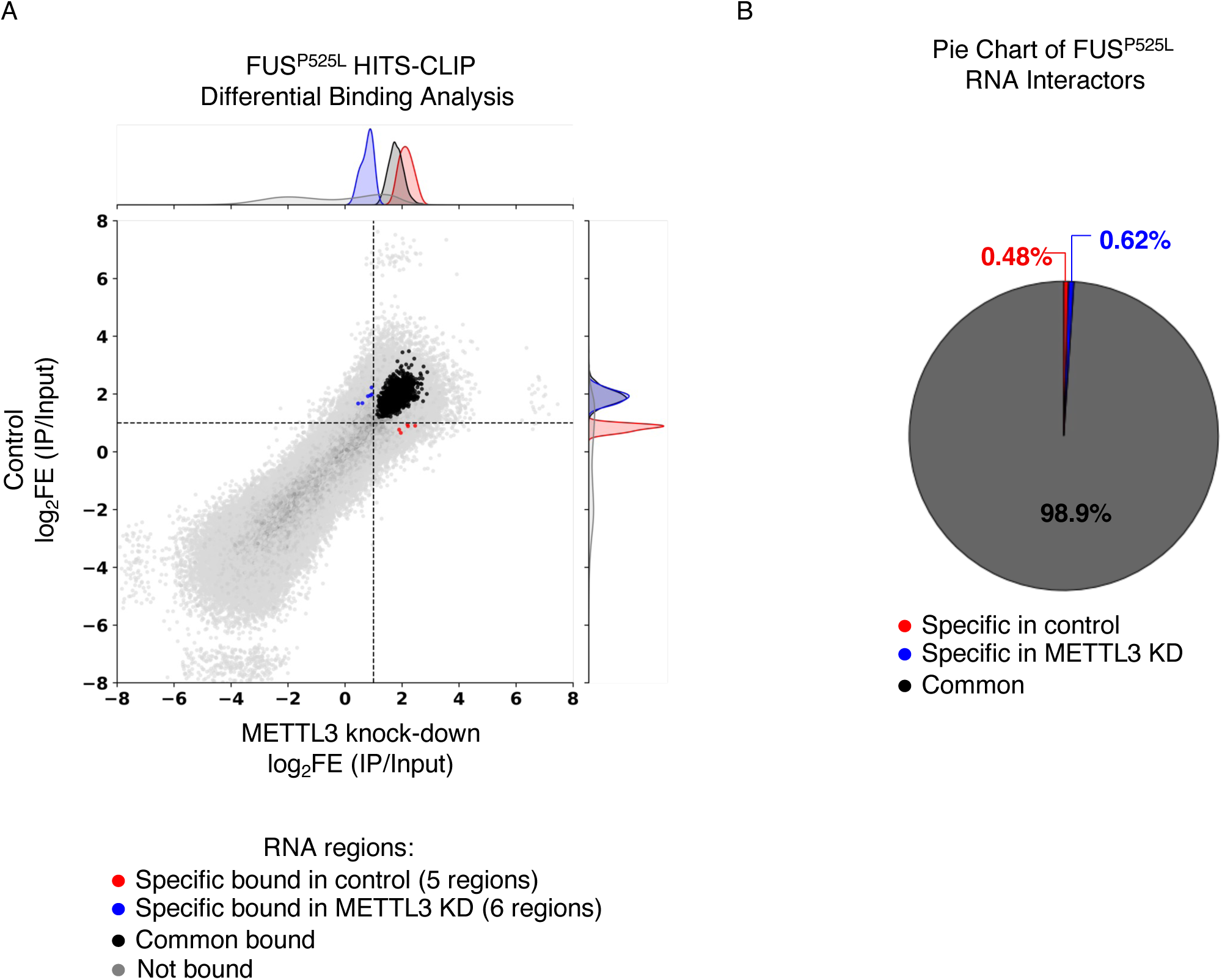
A) Scatter plots show HITS-CLIP log fold-enrichment (log₂FE IP/Input) of FUS^P525L^ binding sites comparing control and METTL3-depleted (mAID) in SK-N-BE cells expressiong FUS^P525L^. Each point represents a genomic region (peak) associated with FUS^P525L^ binding (specific or common) or used as negative control (not bound). Gray dots indicate invariant peaks (not differentially enriched), while red and blue dots represent peaks significantly enriched in FUS^P525L^ and mAID conditions, respectively. Marginal density plots display the distribution of log₂FE. The intensity of gray points reflects local peak density — darker gray indicates higher point density, as determined by a 2D histogram-based alpha adjustment. *n* = 2 biologically independent replicates. B) Pie chart showing the fraction of FUS^P525L^ interactors common or specific to control and METTL3 knock-down conditions.

These data are coherent with previously published data showing that m^6^A target regions *per se* are not preferred sites for FUS binding^2^.

### Mutant FUS drives ALKBH5 downregulation, increasing m^6^A levels

Building on observations from both cellular and animal models, we sought to elucidate the mechanisms underlying the increase in m^6^A levels in mutant FUS-associated ALS. To this aim, we assessed potential alterations of the main factors regulating m^6^A dynamics. Specifically, we compared their RNA and protein levels in SK-N-BE cells expressing either FLAG-tagged wild-type FUS (FUS^WT^) or the ALS-linked mutant FUS^P525L^ (FUS^P525L^). We also included their respective METTL3 knockdown counterparts, generated by CRISPR-Cas9-mediated insertion of a degron-tag at the N-terminus of METTL3 (mAID-FUS^WT^ and mAID-FUS^P525L^)^2^. As a first step, we assessed the expression of the core m^6^A writer complex components, METTL3 and METTL14. ^27^As shown in Fig. 3A, no differences were detected between FUS^P525L^ and FUS^WT^ cells.

**Figure 3.**
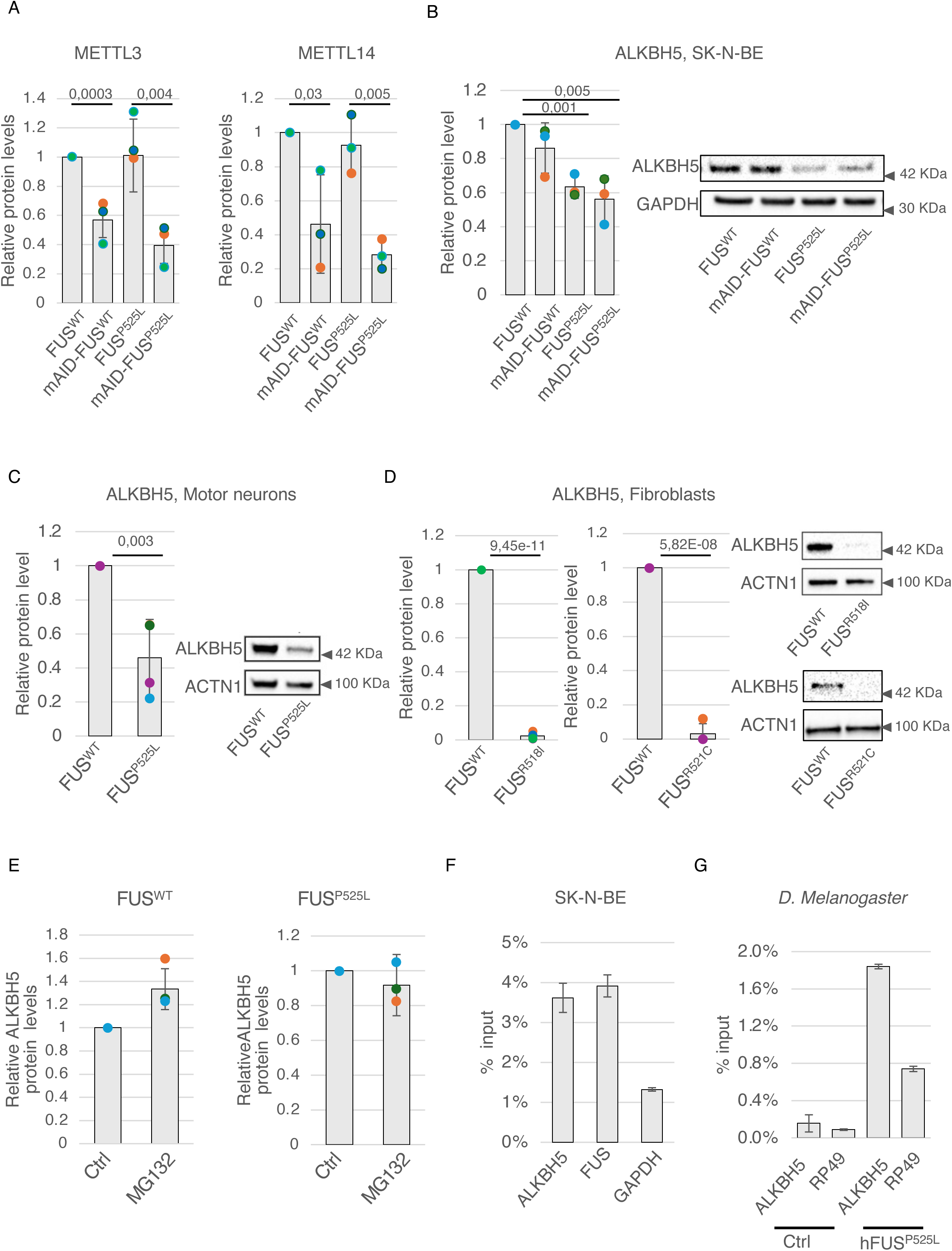
A) Relative protein levels showing the levels of either METTL3 or METTL14 in the indicated SK-N-BE cells. ACT-β or ACTN1 were used as loading control. Relative protein levels were represented as relative quantities with respect to control (FUS^WT^) cells set as 1. The relative protein quantity in the bars is represented as mean of replicates with standard deviation. Dots represent independent replicates. n=4 biologically independent replicates. The *ratio* of each sample *versus* its experimental control was tested by two-tailed Student’s t test. P-values are indicated. B, C, D) Representative western blot analysis (right) and the corresponding densitometry analyses (left) showing the levels of ALKBH5 in the indicated cells. GAPDH or ACTN1 were used as loading control. Relative protein levels were represented as relative quantities with respect to control (FUS^WT^) cells set as 1. The relative protein quantity in the bars is represented as mean of replicates with standard deviation. Dots represent independent replicates. n=3 biologically independent replicates. The *ratio* of each sample *versus* its experimental control was tested by two-tailed Student’s t test. P-values are indicated. E) Densitometric analyses of ALKBH5 protein levels obtained by Western blot analysis, using ALKBH5 antibodies, of proteins from the indicated SK-N-BE cells treated with DMSO (Ctrl) or MG132. ACTN1 was used as loading control. Relative protein levels were represented as relative quantities with respect to control (Ctrl) cells set as 1. The relative protein quantity in the bars is represented as mean of replicates with standard deviation. Dots represent independent replicates. n=3 biologically independent replicates. The *ratio* of each sample *versus* its experimental control was tested by two-tailed Student’s t test. P-values are indicated. F, G) Bar plot showing the enrichment of the ALKBH5 transcript in a representative FUS-CLIP experiment performed either in SK-N-BE (F) cells or *D. Melanogaster* extracts (G). Levels are represented as percentage with respect to the input. n=3 biologically independent replicates. GAPDH or RP49 have been used as negative controls, while FUS has been used as positive control in SK-N-BE cells.

In the mAID lines, METTL3 was efficiently downregulated, accompanied by a parallel decrease in its interactor METTL14, consistent with the stabilizing effect of METTL3 on this protein^28^. We then assessed the m^6^A eraser ALKBH5 and the main cytoplasmic and nuclear readers: YTHDF1, YTHDF2, YTHDF3, and YTHDC1. Moreover, to explore potential deregulation in the metabolism of S-adenosylmethionine (SAM), the methyl-donor required for m^6^A deposition, we also examined METTL16 and MAT2A, both involved in SAM biosynthesis^29,30^.

None of the tested factors exhibited substantial RNA or protein-level deregulation compared to controls (Fig. S3A and B), with the exception of YTHDC1 and ALKBH5. Interestingly, ALKBH5 while showing a modest decrease (∼20%) of the RNA (Fig. S3A) exhibited a pronounced reduction at the protein level (50%) in FUS^P525L^-expressing cells (Fig. 3B). ALKBH5 reduction provides a plausible explanation for the increased m^6^A levels observed in the mutant condition, coherently with the fact that the decrease of this m^6^A eraser provokes m^6^A accumulation^31^. Importantly, reduction of ALKBH5 protein levels was also confirmed in iPSC-derived motor neurons carrying the FUS^P525L^ mutation (Fig. 3C), as well as in patient-derived fibroblasts carrying either the FUS^R518I^ or FUS^R521C^ mutations (Fig. 3D) with respect to their experimental controls. We did not observe any alteration of the ALKBH5 RNA levels in those systems (Fig. S3C, S3D).

YTHDC1, instead, resulted up-regulated in all conditions with respect to the FUS^WT^ cell line (Fig. S3B). Therefore, we considered this change unable to explain the global m^6^A increase observed specifically upon FUS^P525L^ expression.

Notably, in our previous study we demonstrated that RNAi-mediated ALKBH5 downregulation, as well as METTL3 overexpression, were sufficient to reproduce in FUS^WT^ cells the increased number of SG observed in FUS^P525L^ cells ^2^. Consistently, both conditions further exacerbated the aberrant SG phenotype in FUS^P525L^ cells^2^.

Since ALKBH5 mRNA downregulation was modest and inconsistent across cellular models (Fig. S23A, S3C, S3D), yet protein levels were consistently reduced in all of them, we investigated whether impaired translation or enhanced degradation underlie the decreased steady-state amounts of ALKBH5. To assess if reduced ALKBH5 protein levels in FUS^P525L^ cells could be due to enhanced degradation, we first performed a cycloheximide (CHX) chase assay. CHX inhibits translation elongation, allowing evaluation of protein stability independent of new production. If ALKBH5 was more rapidly degraded in the presence of mutant FUS, its levels would be expected to decline more rapidly upon CHX treatment with respect to the non-treated control. Instead, while we observed ALKBH5 protein levels decreasing in FUS^WT^ condition, it remained stable in FUS^P525L^ cells compared to FUS^WT^ controls (Fig. S3E), suggesting that an enhanced protein degradation is unlikely to explain the observed reduction.

As a complementary approach, we carried out proteasome inhibition assays using MG132, a compound that blocks proteasome-mediated protein degradation. If the reduction in ALKBH5 levels were due to proteasome-mediated protein degradation, MG132 treatment should have led to an increase in the ALKBH5 protein. However, our data show no change in ALKBH5 upon proteasome inhibition, further proving that the decrease observed in mutant FUS cells is not caused by enhanced degradation but rather by impaired translational control (Fig. 3E).

Overall, our data support the conclusion that the increased cytoplasmic levels of FUS occurring in FUS^P525L^ cells do not impair ALKBH5 degradation, but rather suggest a defect in its translation, accounting for the observed ALKBH5 downregulation in ALS cellular models. To further validate this hypothesis, we assessed ALKBH5 protein levels in SK-N-BE cells exposed to increasing doxycycline concentrations to induce gradual FUS overexpression. As shown in Fig. S3F, progressive induction of FUS through escalating doxycycline concentrations leads to a corresponding dose-dependent decrease in ALKBH5 protein levels. These data indicate that FUS levels negatively correlate with ALKBH5 translation in a dose-dependent manner.

Consistently with these findings, CLIP-seq^32^ analysis identified a FUS^P525L^ binding site within the 3′ untranslated region (UTR) of the ALKBH5 transcript.

Piranha peak calling detected this peak in both biological replicates, and its presence was further supported by higher IP than input coverage (Fig. S3G). These data indicate a direct interaction between FUS^P525L^ and ALKBH5 mRNA that may contribute to the observed translational control. We validated the CLIP-seq data by performing a FUS-CLIP followed by RT-qPCR in our FUS^P525L^ SK-N-BE cells (Fig. S3H). The data confirmed the specific enrichment of ALKBH5 mRNA in the FUS^P525L^ IP fraction, with the FUS transcript^33^ and GAPDH serving as positive and negative controls, respectively (Fig. 3F).

Importantly, we also detected FUS binding to ALKBH5 mRNA in extracts from *Drosophila melanogaster* expressing hFUS^P525L^ (Fig. 3G; S3I). No enrichment of ALKBH5 was detected in immunoprecipitates from wild-type flies, non-expressing hFUS^P525L^, which served as an additional control. This supports the notion that the observed ALKBH5 transcript enrichment specifically occurs in the presence of mutant FUS.

To gather some insights on the FUS^P525L^-mediated translational regulation of ALKBH5, we investigated whether other RNA-binding proteins (RBP), known to affect translation might interact with the same region of the ALKBH5 transcript interacting with FUS. Using our HITS-CLIP data^2^, we confirmed the putative FUS binding site on ALKBH5 mRNA. We then employed an *in silico* screening approach to assess the potential co-binding of other RBP to this region. Specifically, we used motif affinity scoring based on PEKA^34^ scores, POSTAR3 CLIPdb RBPs binding regions^20^ and RNAcompete-derived binding preferences^35^ (see Materials and Methods) to rank RBP by their predicted affinity for the ALKBH5 transcript, specifically at the level of the FUS interacting region. High-confidence RBP (Z-score > 3) enriched at the FUS-binding site were selected (Fig. S3G). Among the top candidates, we identified HUR, TIA1, and PTBP1, three well-characterized RBP known for their roles in RNA translation regulation^36–40^ (Fig. S3G).

The presence of these RBP in proximity of the FUS-binding region suggests a possible regulatory hub, where competitive or cooperative interactions may influence ALKBH5 mRNA fate in response to increased levels of cytoplasmic FUS, such as in the condition of FUS^P525L^ expression. This raises the intriguing possibility that FUS may disrupt or modulate the access of translational regulators on ALKBH5 mRNA, thereby contributing to the altered translational output observed in our system.

In conclusion, our data suggest that ALKBH5 is post-transcriptionally downregulated in mutant FUS-associated ALS cellular models, due to translational repression mediated by FUS, that binds the 3’UTR of ALKBH5 mRNA.

### ALKBH5 downstream targets are altered in mutant FUS-associated ALS and contribute to the SG aberrant phenotype

In order to assess whether the ALKBH5 decrease observed in FUS^P525L^ cells has any downstream target that could impinge on the ALS-related SG phenotype, we tested if selected candidates, altered in mutant FUS conditions, were affected by ALKBH5 RNAi-mediated knockdown.

Candidate selection was based on an integrative analysis of transcriptomic and proteomic datasets. We prioritized genes that showed consistent deregulation in FUS^P525L^ cells compared to FUS^WT^ across multiple datasets, including RNA-seq of SK-N-BE^2^ cells and iPSC-derived motor neurons (MN)^27^, and MN-derived mass spectrometry (MS) data^41^. To capture potentially translationally regulated targets, we also included the top upregulated and downregulated proteins from the MN-MS dataset, regardless of their RNA deregulation in SK-N-BE cells, but excluding transcripts not expressed in this system. The remaining candidates were further refined through a literature-based rational text mining using keywords such as “ALS”, “neurodegeneration”, “stress granules”, and “phase separation”.

This pipeline yielded a shortlist of 18 candidate genes (Table 1). One gene, CHL1, failed to amplify by RT-qPCR and was excluded from subsequent analyses.

**Table 1.**
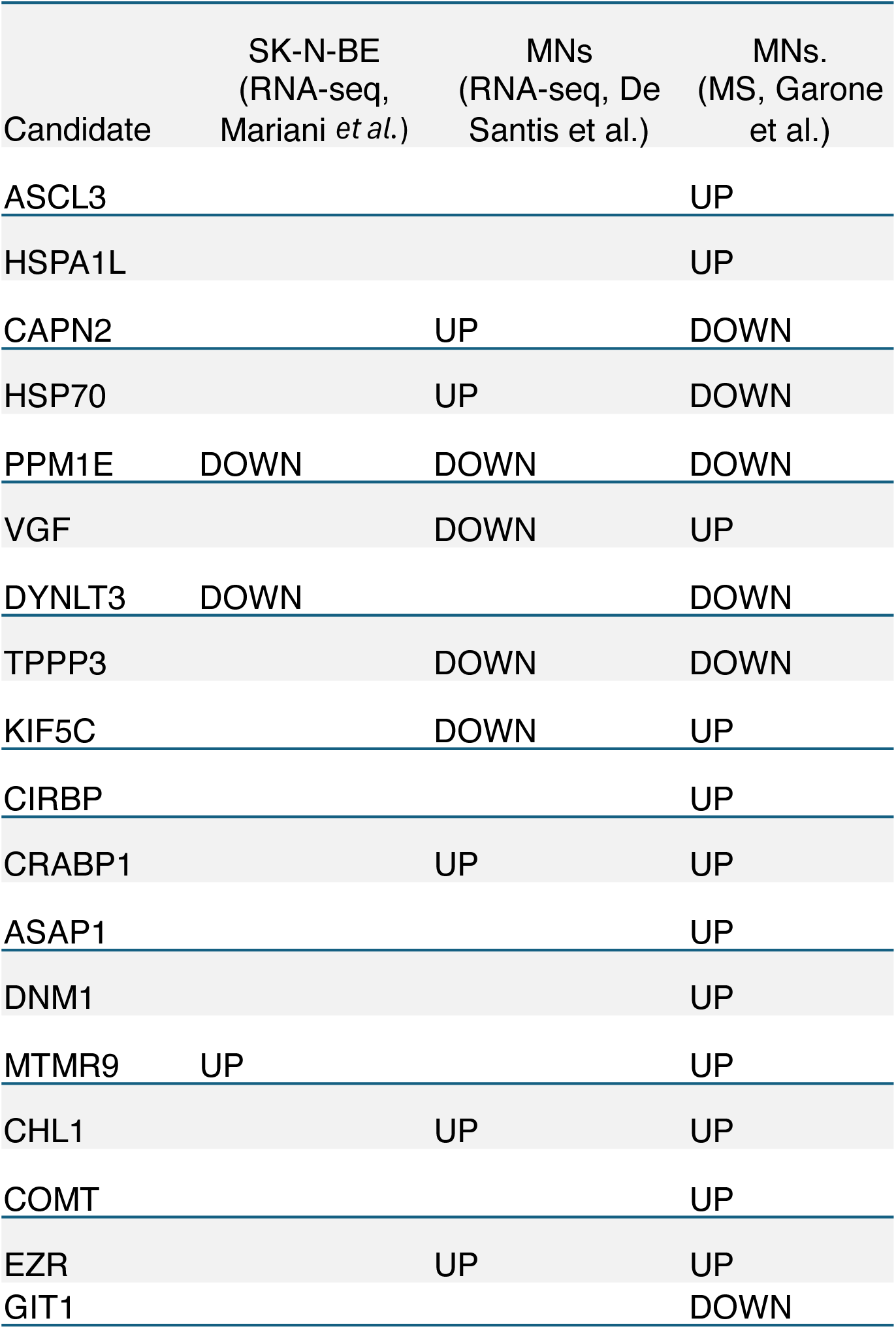
List of the candidates selected basing on an integrative analysis of transcriptomic/proteomic datasets from SK-N-BE and MNs expressing FUS^P525L^ followed by a literature-based rational text mining. Trends of deregulation are specified

To identify transcripts showing consistent deregulation patterns, we depleted ALKBH5 by siRNA in SK-N-BE cells expressing either FUS^WT^ or FUS^P525L^ and profiled the 17 candidates for their responsiveness to ALKBH5 and their deregulation in ALS conditions (Fig. 4A, S4A, S4B).

**Figure 4.**
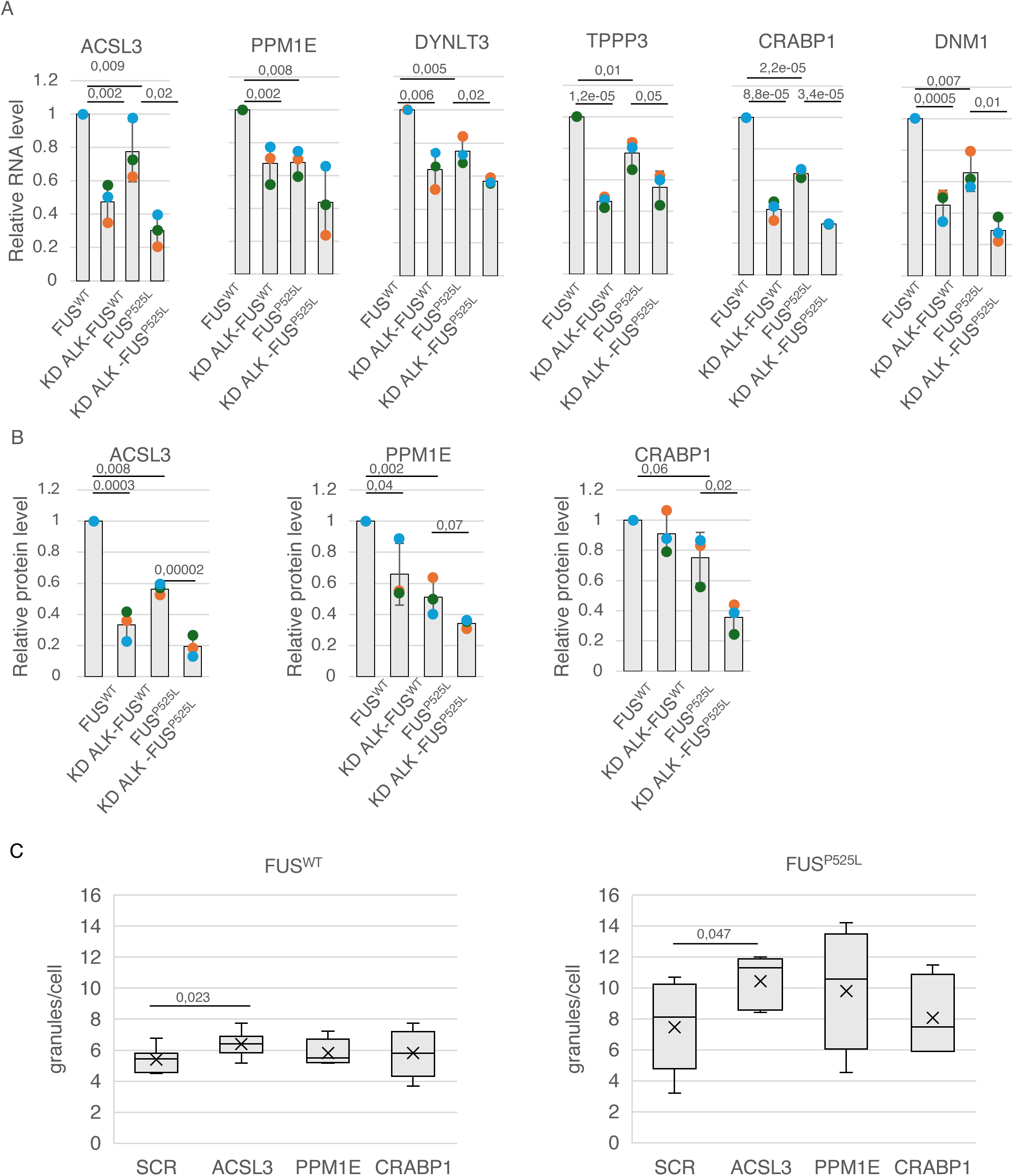
A)Bar plot showing the relative RNA levels of selected candidates in the indicated SK-N-BE cells. Values are normalized against GAPDH transcript and expressed as relative quantity with respect to FUS^WT^ cells set as 1. The relative RNA quantity in the bars is represented as mean of the fold change with standard deviation. Dots represent each replicate. n=3 biologically independent replicates. The *ratio* of each sample *versus* its experimental control was tested by two-tailed Student’s t test. P-values are indicated. B) Densitometry analyses showing the levels of selected candidates in the indicated SK-N-BE cells. ACT-β was used as loading control. Relative protein levels were represented as relative quantities with respect to control (FUS^WT^) cells set as 1. TA) Box plots illustrating the number of stress granules per cell in the indicated stressed SK-N-BE upon si-RNA treatment against selected candidates. The box plots are defined by minima, 25% percentile, median, 75% percentile, and maximum (See Source Data file for values). The x indicates the average number of granules in each sample. 3-5 fields were acquired for each biological replicate. n=3 biologically independent replicates. The *ratio* of each sample *versus* its experimental control (SCR) was tested by two-tailed Student’s t test. P-values are indicated. he relative protein quantity in the bars is represented as mean of replicates with standard deviation. Dots represent independent replicates. n=3 biologically independent replicates. The *ratio* of each sample *versus* its experimental control was tested by two-tailed Student’s t test. P-values are indicated. C) Box plots illustrating the number of stress granules per cell in the indicated stressed SK-N-BE upon si-RNA treatment against selected candidates. The box plots are defined by minima, 25% percentile, median, 75% percentile, and maximum (See Source Data file for values). The x indicates the average number of granules in each sample. 3-5 fields were acquired for each biological replicate. n=3 biologically independent replicates. The *ratio* of each sample *versus* its experimental control (SCR) was tested by two-tailed Student’s t test. P-values are indicated.

Among them, six were downregulated in FUS^P525L^ conditions and upon ALKBH5 knockdown in both FUS^WT^ and FUS^P525L^ cell lines (ACSL3, PPM1E, DYNLT3, TPPP3, CRABP1, DNM1, Fig. 4A).

Protein analysis revealed that three of these genes (ACSL3, PPM1E, CRABP1) displayed also consistent protein downregulation upon ALKBH5 reduction in both FUS^WT^ and FUS^P525L^ cells (Fig.4B). Moreover, we found that the levels of ACSL3, PPM1E, CRABP1 increased when knocking down METTL3 in FUS^P525L^ cells (Fig. S4C), thus reversing ALKBH5 downregulation. Collectively, our data indicate that there is a subset of genes i) downregulated in FUS^P525L^ conditions, ii) specifically responding to ALKBH5 reduction, and iii) rescued when m^6^A levels are restored by METTL3 knockdown. These data support the idea that the alteration of these genes observed in FUS^P525L^ conditions may be m^6^A-dependent.

While for CRABP1 the effect might be indirect since it has been shown not to contain m^6^A modifications according to an m^6^A CLIP experiment (Fig.S4D), for ACSL3 and PPM1E the effect is likely to be direct since these mRNAs are enriched in an m^6^A CLIP experiment, indicating that they are decorated with m^6^A (Fig. S4D).

We previously reported that expression of FUS^P525L^ increases the number of SG per cell, a phenotype that is reversible upon decrease of m^6^A levels by METTL3 knockdown^2^. To test whether the identified ALKBH5 downstream targets may contribute to the SG phenotype, we performed RNAi experiments in SK-N-BE cells expressing either FUS^WT^ or FUS^P525L^ and quantified the number of SG per cell upon arsenite treatment (Fig. 4C, S4F).

Following this *criterium*, we tested the effects of knocking down ACSL3, PPM1E, and CRABP1. Interestingly, the knockdown of ACSL3 and CRABP1 led to a significant SG increase in FUS^WT^ (Fig. 4C). Interestingly, we found that ACSL3 downregulation not only was sufficient to produce SG increase in FUS^WT^ cells but also to worsen such phenotype in mutant cells (Fig. 4C).

Notably, ACSL3 knock-down did not provoke relevant deregulations of ALKBH5, suggesting that it might act downstream of ALKBH5(Fig. S4G).

## Discussion

In previous studies, we established a link between increased m^6^A levels and the aberrant SG dynamics observed in mutant FUS-associated ALS cellular models, supporting the concept of m^6^A as a druggable therapeutic target in ALS.

In this study, on one hand we aimed to strengthen our data with *in vivo* validation in flies; on the other, we investigated the upstream molecular mechanisms driving the increase in m^6^A levels and its downstream consequences.

Here we provide *in vivo* validation of the beneficial effects of m^6^A downregulation in ALS. Indeed, we showed that lowering m^6^A levels (*via* IME4/METTL3 knockdown) alleviates locomotor defects and NMJ abnormalities in Drosophila expressing hFUS^P525L^, demonstrating that modulating m^6^A is not only effective in cell culture, but can also mitigate ALS phenotypes in an entire organism. These results reinforce the idea that restoring m^6^A homeostasis may have therapeutic relevance and represent a crucial step toward preclinical validation.

Concerning the molecular mechanism, we found that reducing m^6^A by METTL3 depletion has only minor effects on the RNA-binding landscape of mutant FUS, pointing out that m^6^A primarily acts downstream of FUS rather than by redirecting its interactome.

Notably, we showed that the global increase in m^6^A observed in FUS-associated ALS models correlates with a consistent reduction of the m^6^A eraser ALKBH5.

Our data converge on a simple axis: mutant FUS accumulates in the cytoplasm and represses translation of the m^6^A eraser ALKBH5 by binding its 3′ UTR; reduced ALKBH5 raises cellular m^6^A level, which in turn remodels the expression of a set of transcripts involved in SG dynamics. This framework not only rationalizes the global increase of m^6^A observed in FUS-ALS models but also identifies intervention points upstream and downstream of such m^6^A alteration.

The dose-dependent inverse relationship between cytoplasmic FUS and ALKBH5 protein, together with FUS binding to the ALKBH5 3′ UTR, suggests a model in which mutant FUS occludes or perturbs regulatory features of this UTR, potentially by competing with or mis-recruiting translational RBPs (for example ELAVL1/HuR, TIA1, PTBP1). Notably, the ALKBH5 transcript is bound by FUS both in human cellular models (SK-N-BE cells, MNs and fibroblasts) and in *D. melanogaster* expressing mutant FUS^P525L^. These data are also in line with previous studies showing that the binding of mutant FUS to specific target mRNAs can alter their translation ^41^.

As mentioned, the reduction of ALKBH5 leads to an increase in m^6^A levels which can affect the fate of specific transcripts. Among the ALKBH5-responsive transcripts we identified, we finally focused on ACSL3. ACSL3 knockdown in FUS^WT^ cells phenocopies the ALS-like SG number alterations and further exacerbates SG accumulation in FUS^P525L^ cells without altering ALKBH5 itself. These results place ACSL3 as a downstream effector of ALKBH5 and m^6^A upregulation. It is important to note that METTL3 depletion, which restores m^6^A balance and SG dynamics, rescues ACSL3 deregulation in the mutant context.

Importantly, ACSL3 has been shown to be downregulated in Alzheimer’s disease mouse models and to promote BDNF and VEGF-C signalling pathways. Thus, ACSL3 has been highlighted as neuroprotective in the brain and it has been proposed as a therapeutic target for alleviating anxiety and depression associated with such disease^42^.

It is important to note the first-in-class clinical small molecule inhibitor of METTL3, STC-15, showed promising interim Phase 1/2 results against advanced malignancies, offering the possibility for STC-15 to be repurposed for FUS-associated ALS^43–45^ .

However, while global METTL3 inhibition is effective experimentally, global manipulation of m^6^A may raise issues of specificity and safety: m^6^A is ubiquitous and involved in numerous physiological processes. Concerning this point, our mechanistic insights carry therapeutic consequences. Indeed, firstly, our data suggests that, in FUS-ALS, intervention could be aimed at restoring physiological levels altered by a specific defect (ALKBH5 downregulation). Secondly, we linked m^6^A alteration with notable molecular targets that could be exploited in therapy.

Looking ahead, more refined strategies, such as restoring ALKBH5 activity where it is pathologically repressed; disrupting the pathological FUS-ALKBH5 mRNA interaction with decoy oligonucleotides or small molecules; and targeting downstream effectors such as ACSL3 could offer a more favourable therapeutic profile than downregulating the entire writer complex.

## Data availability

The data supporting the findings of this study are available from the corresponding authors upon request. The high throughput sequencing data generated in this study have been deposited in the GEO database. The following secure token has been created to allow review of record GSE310551 while it remains in private status: ifejgqeyllmnhyr.

## Code availability

All software, links to websites, or tools used for this work are referred to in the methods section or in the figure legends. Additional dedicated scripts developed for this work are available upon request.

## Contributions

G.D.T., and I.B. designed and conceived the study. The experiments were performed and analyzed by G.D.T., F.M., A.G., M.C.B., T.S.. Bioinformatics data analysis was performed by A.S. and A.D. Experiments performed in *D. Melanogaster* were performed by M.D.S. and G.C. The original draft of the manuscript was written by I.B. and G.D.T. with suggestions from all the other authors. I.B. supervised the project.

## Acknowledgements

We thank M. Marchioni and V. De Turris for technical help and M. Caruso for assistance. We thank M. Morlando, A. Fatica, A. Rosa, M. Ballarino for discussion.

This work was partially supported by grants from ERC-2019-SyG 855923-ASTRA, AIRC IG 2019 Id. 23053, and PRIN 2017 2017P352Z4 to I.B.; “National Center for Gene Therapy and Drug based on RNA Technology” (CN00000041) and NextGenerationEU PNRR MUR to I.B.; Sapienza departmental projects 2023 (RD12318A998C70B8) to G.D.T.

