## Supplementary Figures S1-4 for "Mutant FUS perturbs m^6^A regulation by repressing ALKBH5, while restoring m^6^A levels alleviates ALS pathology *in vivo*"

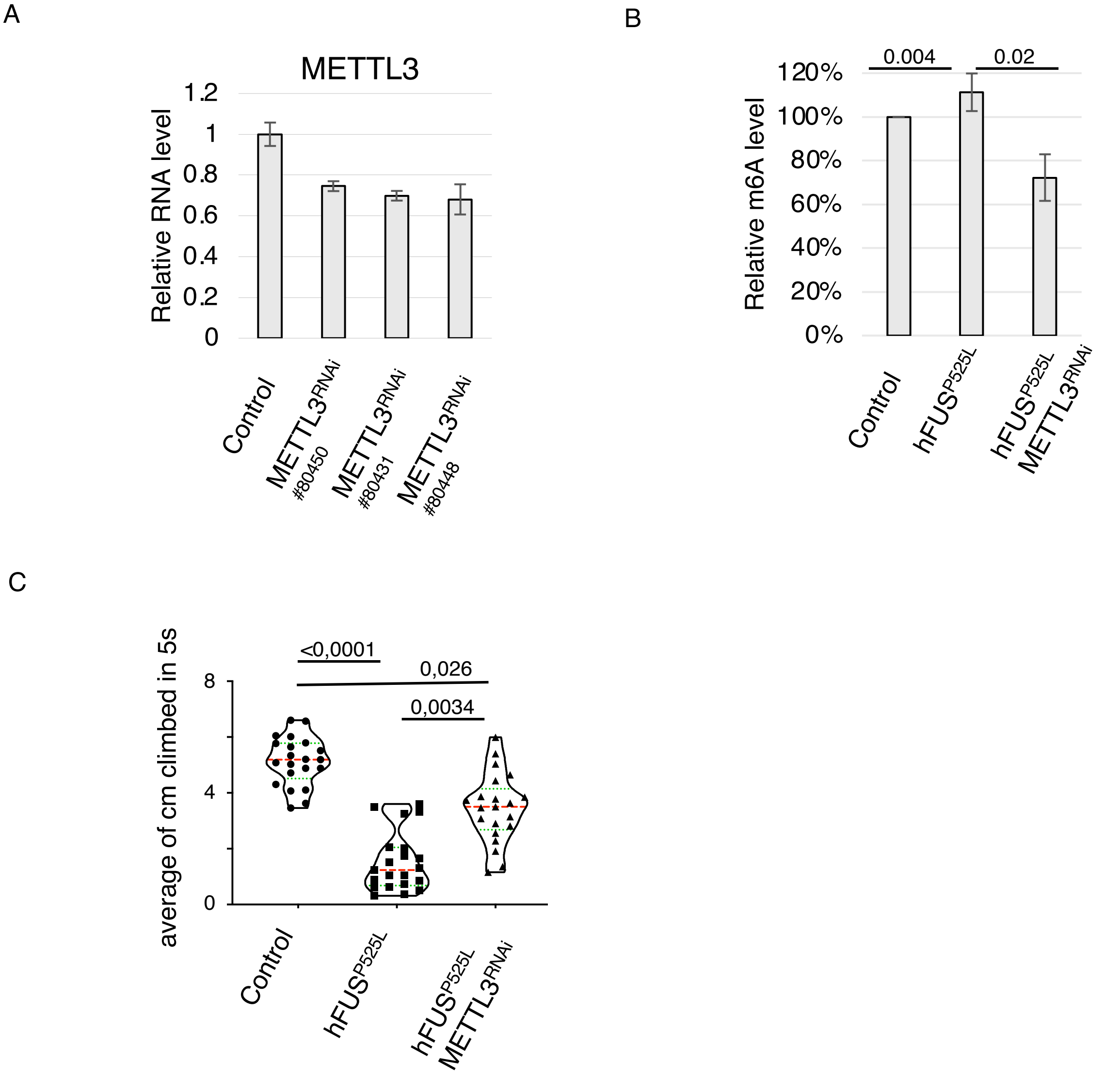


**Figure S1**

A) Bar plot showing METTL3 relative RNA level decrease in the indicated fly strains. Values are normalized against GIOTTO transcript and expressed as relative quantity with respect to control 69B-GAL4 flies set to a value of 1. The relative RNA quantity in the bars is represented as mean of the fold change with standard deviation. B) Bar plot showing the m$^6$A level in the indicated fly strains. Levels are represented as percentage with respect to control flies set as 100%. n=2 biologically replicates. The relative m$^6$A percentage in the bars is represented as mean of the replicates with standard deviation. The ratio of each sample versus its experimental control was tested by two-tailed Student's t test. P-values are indicated. C) Quantification of fly locomotor activity assessing both the average distance climbed by adult male flies expressing the indicated RNAi constructs or transgenes, under control of 69B-GAL4 in 5 seconds (control n=76 flies, hFUS$^{P525L}$ n=77 flies, hFUS$^{P525L}$METTL3$^{RNAi}$ n=79 flies and the average distance (cm) climbed (right panel). The truncated violin plot reports median (dashed lines), first and third quartile (dotted lines) and density plot (outside lines). One way ANOVA and Tukey's multiple comparison tests were used to compare samples, p-value are indicated.

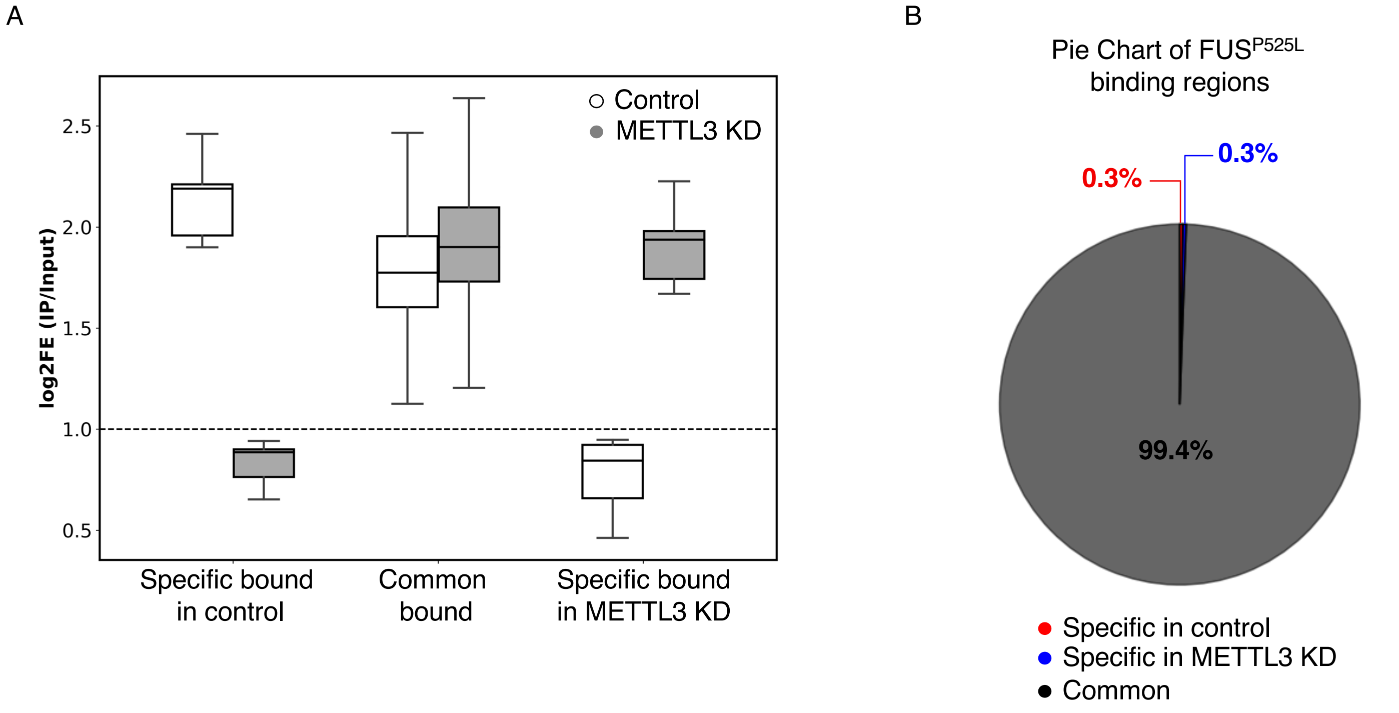


**Figure S2**

A) Boxplot showing the HITS-CLIP $log_2$ fold-enrichment (IP/Input) of $FUS^{P525L}$ binding sites in SK-N-BE cells, comparing control and METTL3 KD (mAID) conditions across control-specific, common, and mAID-specific regions. B) Pie chart showing the fraction of $FUS^{P525L}$ binding regions common or specific to control and METTL3 KD conditions.

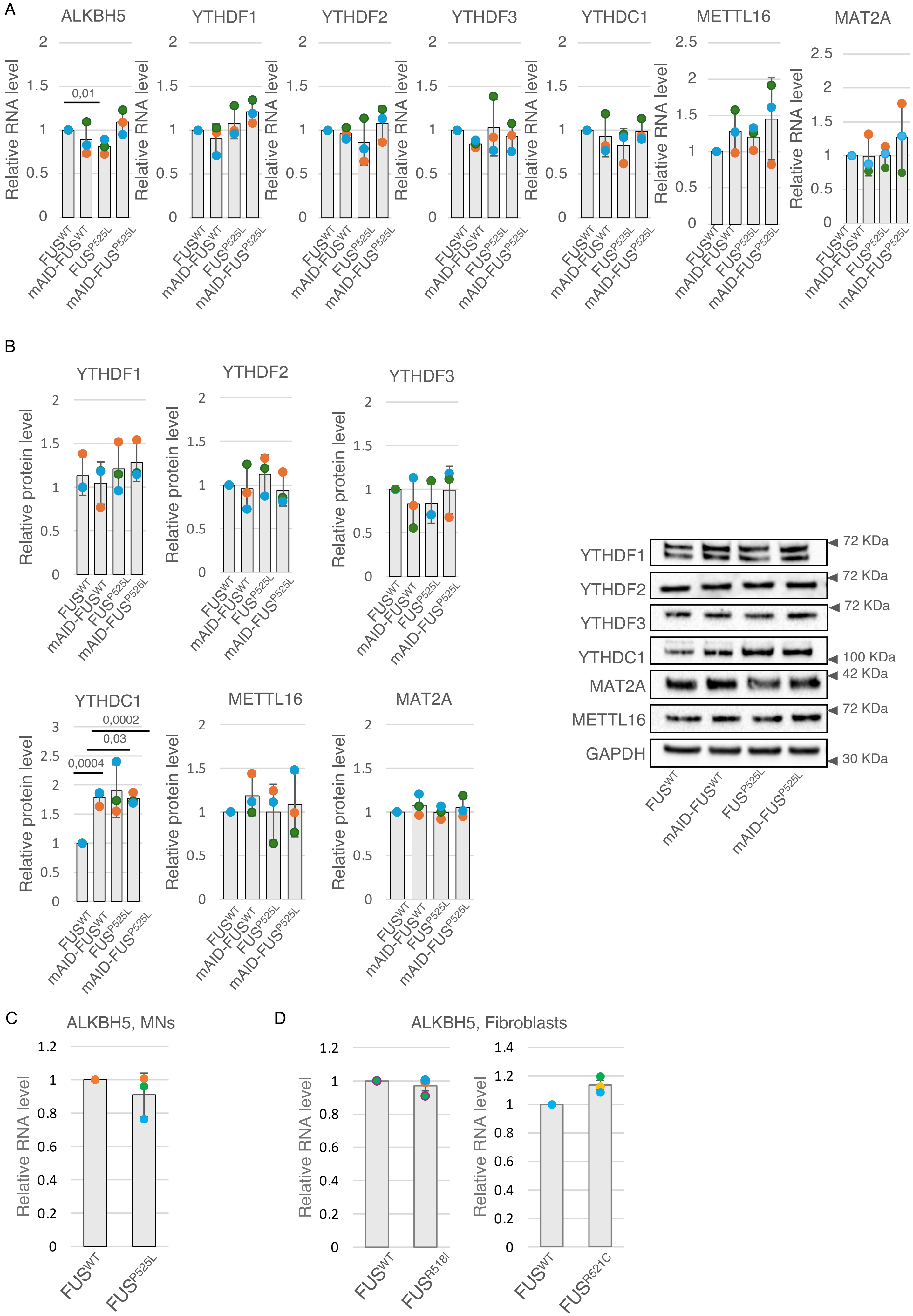

A
ALKBH5
YTHDF1
YTHDF2
YTHDF3
YTHDC1
METTL16
MAT2A
Relative RNA level
0,01
FUS^WT
mAID-FUS^WT
FUS^P525L
mAID-FUS^P525L
B
YTHDF1
YTHDF2
YTHDF3
YTHDC1
METTL16
MAT2A
Relative protein level
0,0002
0,03
0,0004
YTHDF1
YTHDF2
YTHDF3
YTHDC1
MAT2A
METTL16
GAPDH
72 KDa
72 KDa
72 KDa
100 KDa
42 KDa
72 KDa
30 KDa
FUS^WT
mAID-FUS^WT
FUS^P525L
mAID-FUS^P525L
C
ALKBH5, MNs
Relative RNA level
FUS^WT
FUS^P525L
D
ALKBH5, Fibroblasts
Relative RNA level
FUS^WT
FUS^R518I
FUS^WT
FUS^R521C

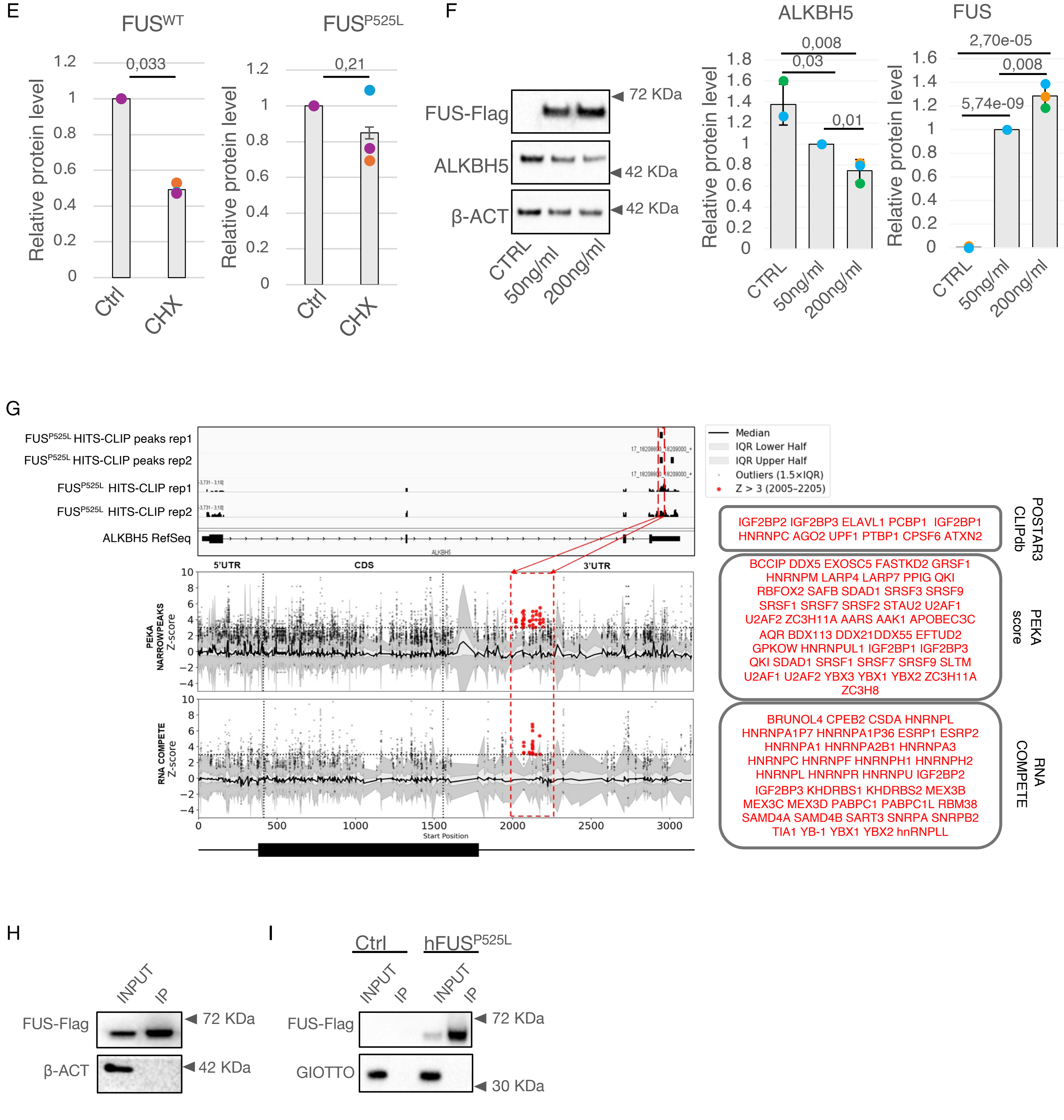


**Figure S3**

A) Bar plot showing the relative RNA levels of selected m$^6$A-related factors in the indicated SK-N-BE cells. Values are normalized against GAPDH transcript and expressed as relative quantity with respect to FUS$^{WT}$ cells set as 1. The relative RNA quantity in the bars is represented as mean of the fold change with standard deviation. Dots represent each replicate. n=3 biologically independent replicates. The *ratio* of each sample *versus* its experimental control was tested by two-tailed Student's t test. P-values are indicated. B) Representative western blot analysis (right) and the corresponding densitometry analyses (left) showing the levels of selected m$^6$A-related factors in the indicated SK-N-BE cells. GAPDH was used as loading control. Relative protein levels were represented as relative quantities with respect to control (FUS$^{WT}$) cells set as 1. The relative protein quantity in the bars is represented as mean of replicates with standard deviation. Dots represent independent replicates. n=3 biologically independent replicates. The *ratio* of each sample *versus* its experimental control was tested by two-tailed Student's t test. P-values are indicated. C, D) Bar plot showing the relative RNA levels of ALKBH5 in the indicated motor neurons and fibroblasts cells. Values are normalized against GAPDH transcript and expressed as relative quantity with respect to FUS$^{WT}$ cells set as 1. The relative RNA quantity in the bars is represented as mean of the fold change with standard deviation. Dots represent each replicate. n=3 biologically independent replicates. The *ratio* of each sample *versus* its experimental control was tested by two-tailed Student's t test. P-values are indicated.

E) Densitometry analyses showing the levels of ALKBH5 in the indicated SK-N-BE cells treated or not with CHX. ACT-β was used as loading control. Relative protein levels were represented as relative quantities with respect to control sample (“Ctrl”) set as 1. The relative protein quantity in the bars is represented as mean of replicates with standard deviation. Dots represent independent replicates. n=3 biologically independent replicates. The *ratio* of each sample *versus* its experimental control was tested by two-tailed Student’s t test. P-values are indicated. F) Representative western blot analysis (left) and the corresponding densitometry analyses (right) showing the levels of either ALKBH5 or FUS in $FUS^{P525L}$ SK-N-BE cells treated with either different doses of doxycicline (50 or 200 ng/ml) or with DMSO, used as control. ACT-β was used as loading control. Relative protein levels were represented as relative quantities with respect to control (CTRL) cells set as 1. The relative protein quantity in the bars is represented as mean of replicates with standard deviation. Dots represent independent replicates. n=3 biologically independent replicates. The *ratio* of each sample *versus* its experimental control was tested by two-tailed Student’s t test. P-values are indicated. G) Left panel: RBP motif-enrichment profiles along the ALKBH5 transcript, based on PEKA  and RNAcompete ( z-scores. The black curve represents the median z-score at each nucleotide position, with shaded areas indicating the interquartile range (IQR): light gray for the central 50% (25th–75th percentile) and darker gray for the extended range defined by 1.5×IQR. Protein outliers whose motif-enrichment scores fall outside this range are shown as gray dots. High-confidence RBPs — defined as proteins with z-scores > 3 within the $FUS^{P525L}$ binding region (positions 2005–2205) — are represented as red dots. Vertical dashed lines mark the boundaries of the 5′UTR, coding sequence (CDS), and 3′UTR, while the red dashed box highlights the $FUS^{P525L}$ binding region. Right panel: Heatmaps showing z-scores for individual RBPs with significant motif enrichment (z > 3) within the $FUS^{P525L}$ binding region, based on POSTAR3 CLIPdb, PEKA, and RNAcompete. Rows represent RBPs, and columns correspond to 5-nucleotide windows spanning positions 2005–2205 of the ALKBH5 transcript. Color intensity reflects the motif-enrichment strength at each position within the FUS binding hotspot. A boxed list on the side reports the RBPs with POSTAR3 CLIPdb-supported binding sites within this region. H) I) Western blot analyses showing the enrichment of the FUS in a representative FUS-CLIP experiment performed either in SK-N-BE cells (H) or *D. Melanogaster* extracts (I). ACT-β or GIOTTO have been used as negative control.

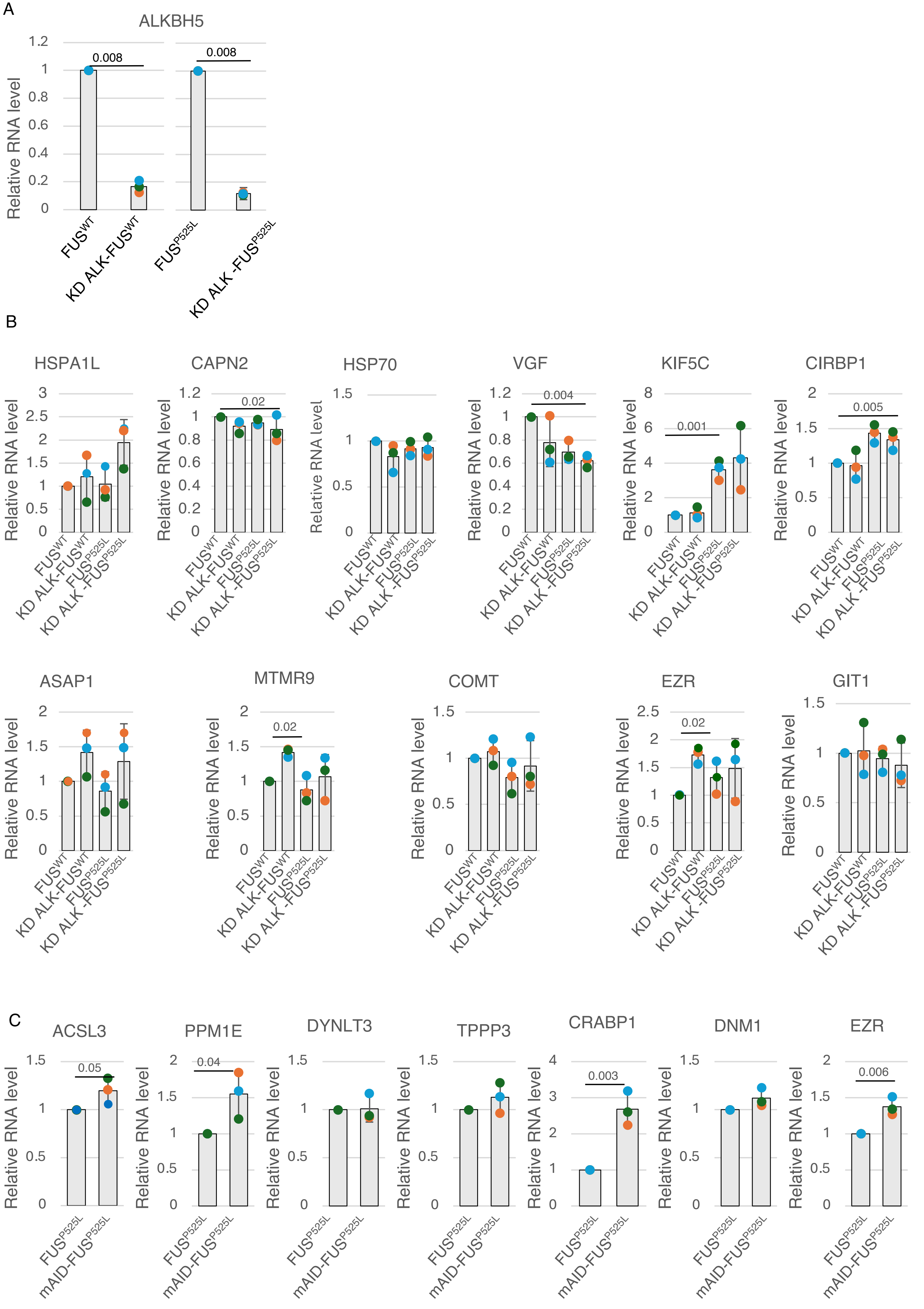

A
ALKBH5
Relative RNA level
0.008
0.008
FUS^WT
KD ALK-FUS^WT
FUS^P525L
KD ALK -FUS^P525L
B
HSPA1L
CAPN2
HSP70
VGF
KIF5C
CIRBP1
0.02
0.004
0.001
0.005
ASAP1
MTMR9
COMT
EZR
GIT1
0.02
0.02
Relative RNA level
FUS^WT
KD ALK-FUS^WT
FUS^P525L
KD ALK -FUS^P525L
C
ACSL3
PPM1E
DYNLT3
TPPP3
CRABP1
DNM1
EZR
0.05
0.04
0.003
0.006
Relative RNA level
FUS^P525L
mAID-FUS^P525L

D

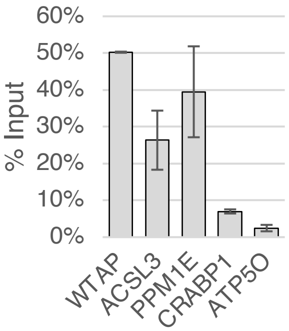
% Input
60%
50%
40%
30%
20%
10%
0%
WTAP
ACSL3
PPM1E
CRABP1
ATP5O

E

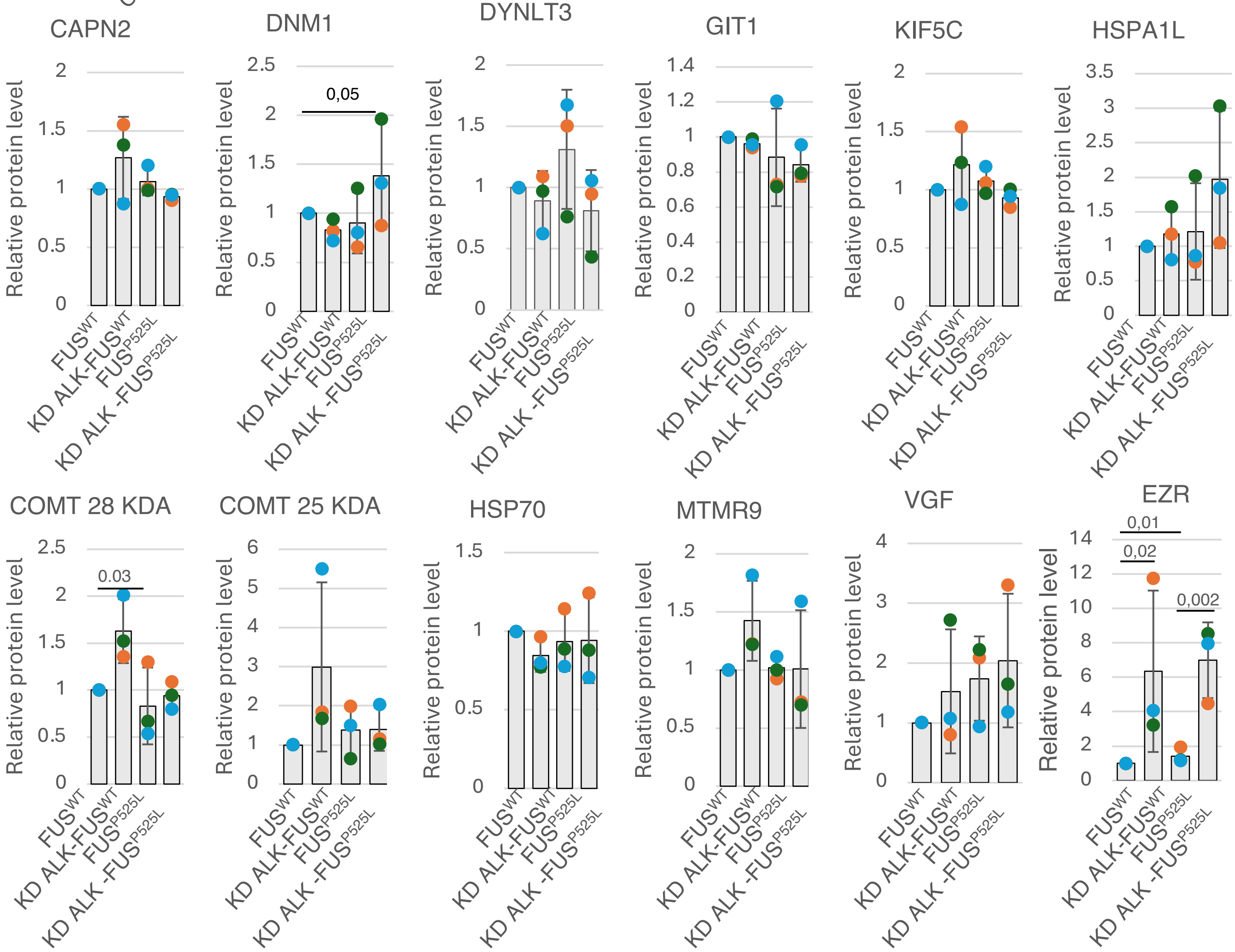
CAPN2
DNM1
DYNLT3
GIT1
KIF5C
HSPA1L
COMT 28 KDA
COMT 25 KDA
HSP70
MTMR9
VGF
EZR
Relative protein level
0,05
0.03
0,01
0,02
0,002
FUSWT
KD ALK-FUSWT
FUSP525L
KD ALK -FUSP525L

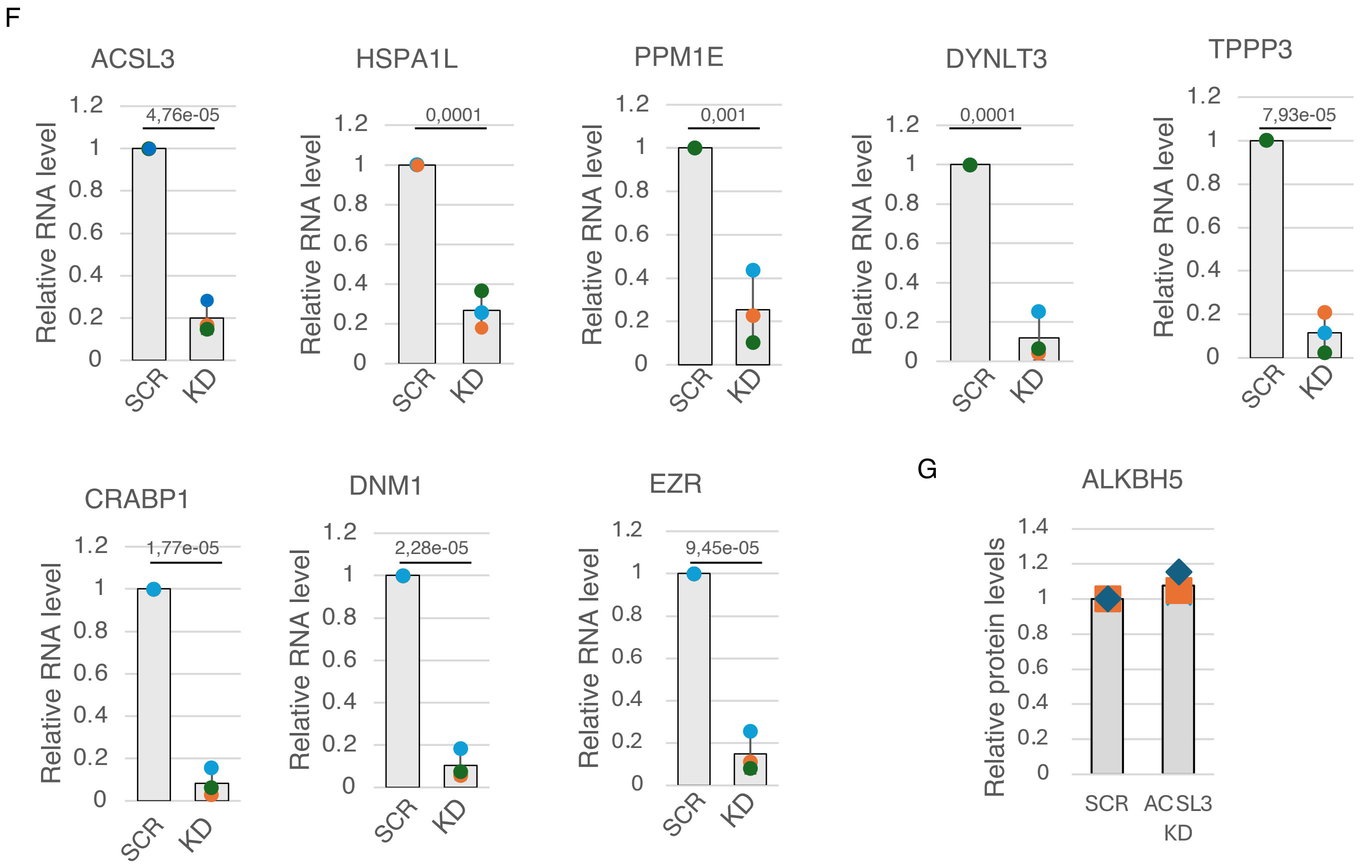


**Figure S4**

A) Bar plot showing the relative RNA levels of ALKBH5 in the indicated SK-N-BE cells in control and ALKBH5 si-RNA condition. Values are normalized against GAPDH transcript and expressed as relative quantity with respect to control cells set as 1. The relative RNA quantity in the bars is represented as mean of the fold change with standard deviation. Dots represent each replicate. n=3 biologically independent replicates. The *ratio* of each sample *versus* its experimental control was tested by two-tailed Student's t test. P-values are indicated. B) Bar plot showing the relative RNA levels of selected candidates in the indicated SK-N-BE cells. Values are normalized against GAPDH transcript and expressed as relative quantity with respect to control cells ($FUS^{WT}$) set as 1. The relative RNA quantity in the bars is represented as mean of the fold change with standard deviation. Dots represent each replicate. n=3 biologically independent replicates. The *ratio* of each sample *versus* its experimental control was tested by two-tailed Student's t test. P-values are indicated. C) Bar plot showing the relative RNA levels of selected candidates in $FUS^{P525}$ SK-N-BE cells either in control or ALKBH5 knock-down condition. Values are normalized against GAPDH transcript and expressed as relative quantity with respect to control cells ($FUS^{WT}$) set as 1. The relative RNA quantity in the bars is represented as mean of the fold change with standard deviation. Dots represent each replicate. n=3 biologically independent replicates. The *ratio* of each sample *versus* its experimental control was tested by two-tailed Student's t test. P-values are indicated. D) Bar plots showing the individual enrichment reduction of selected candidates in a representative m6A CLIP experiment in $FUS^{P525}$ SK-N-BE cells. WTAP and ATP5O were used as positive and negative control, respectively. E) Densitometry analyses showing the levels of selected candidates in the indicated SK-N-BE cells. ACT-β was used as loading control. Relative protein levels were represented as relative quantities with respect to control ($FUS^{WT}$) cells set as 1. The relative protein quantity in the bars is represented as mean of replicates with standard deviation. Dots represent independent replicates. n=3 biologically independent replicates. The *ratio* of each sample *versus* its experimental control was tested by two-tailed Student's t test. P-values are indicated. F) Bar plot showing the relative RNA levels of selected candidates in the indicated SK-N-BE cells in control condition and upon their knock-down. Values are normalized against GAPDH transcript and expressed as relative quantity with respect to control cells ($FUS^{WT}$) set as 1. The relative RNA quantity in the bars is represented as mean of the fold change with standard deviation. Dots represent each replicate. n=3 biologically independent replicates. The *ratio* of each sample *versus* its experimental control was tested by two-tailed Student's t test. P-values are indicated. G) Densitometry analyses showing the levels of ALKBH5 in $FUS^{P525}$ SK-N-BE cells either in control or the indicated knock-down conditions. ACT-β was used as loading control. Relative protein levels were represented as relative quantities with respect to control (SCR) cells set as 1. The relative protein quantity in the bars is represented as mean of replicates with standard deviation. Dots represent independent replicates. n=3 biologically independent replicates. The *ratio* of each sample *versus* its experimental control was tested by two-tailed Student's t test. P-values are indicated.

| Candidate | SK-N-BE (RNA-seq, Mariani *et al.*) | MNs (RNA-seq, De Santis et al.) | MNs. (MS, Garone et al.) |
|---|---|---|---|
| ASCL3 | | | UP |
| HSPA1L | | | UP |
| CAPN2 | | UP | DOWN |
| HSP70 | | UP | DOWN |
| PPM1E | DOWN | DOWN | DOWN |
| VGF | | DOWN | UP |
| DYNLT3 | DOWN | | DOWN |
| TPPP3 | | DOWN | DOWN |
| KIF5C | | DOWN | UP |
| CIRBP | | | UP |
| CRABP1 | | UP | UP |
| ASAP1 | | | UP |
| DNM1 | | | UP |
| MTMR9 | UP | | UP |
| CHL1 | | UP | UP |
| COMT | | | UP |
| EZR | | UP | UP |
| GIT1 | | | DOWN |

**Table 1**
List of the candidates selected basing on an integrative analysis of transcriptomic/proteomic datasets from SK-N-BE and MNs expressing $FUS^{P525L}$ followed by a literature-based rational text mining. Trends of deregulation are specified
